# Elucidation of the interactions between SARS-CoV-2 Spike protein and wild and mutant types of IFITM proteins by *in silico* methods

**DOI:** 10.1101/2021.09.13.460130

**Authors:** Nazli Irmak Giritlioglu, Gizem Koprululu Kucuk

## Abstract

COVID-19 is a viral disease that has been a threat to the whole world since 2019. Although effective vaccines against the disease have been developed, there are still points to be clarified about the mechanism of SARS-CoV-2, which is the causative agent of COVID-19. In this study, we determined the binding energies and the bond types of complexes formed by open (6VYB) and closed (6VXX) forms of the Spike protein of SARS-CoV-2 and wild and mutant forms of IFITM1, IFITM2, and IFITM3 proteins using the molecular docking approach. First, all missense SNPs were found in the NCBI Single Nucleotide Polymorphism database (dbSNP) for IFITM1, IFITM2, and IFITM3 and analyzed with SIFT, PROVEAN, PolyPhen-2, SNAP2, Mutation Assessor, and PANTHER cSNP web-based tools to determine their pathogenicity. When at least four of these analysis tools showed that the SNP had a pathogenic effect on the protein product, this SNP was saved for further analysis. Delta delta G (DDG) and protein stability analysis for amino acid changes were performed in the web-based tools I-Mutant, MUpro, and SAAFEC-SEQ. The structural effect of amino acid change on the protein product was made using the HOPE web-based tool. HawkDock server was used for molecular docking and Molecular Mechanics/Generalized Born Surface Area (MM/GBSA) analysis and binding energies of all complexes were calculated. BIOVIA Discovery Studio program was utilized to visualize the complexes. Hydrogen bonds, salt bridges, and non-bonded contacts between Spike and IFITM protein chains in the complexes were detected with the PDBsum web-based tool. The best binding energy among the 6VYB-IFITM wild protein complexes belong to 6VYB-IFITM1 (-46.16 kcal/mol). Likewise, among the 6VXX-IFITM wild protein complexes, the most negative binding energy belongs to 6VXX-IFITM1 (-52.42 kcal/mol). An interesting result found in the study is the presence of hydrogen bonds between the cytoplasmic domain of the IFITM1 wild protein and the S2 domain of 6VYB. Among the Spike-IFITM mutant protein complexes, the best binding energy belongs to the 6VXX-IFITM2 N63S complex (-50.77 kcal/mol) and the worst binding energy belongs to the 6VXX-IFITM3 S50T complex (4.86 kcal/mol).

The study suggests that IFITM1 protein may act as a receptor for SARS-CoV-2 Spike protein. Assays must be advanced from *in silico* to *in vitro* for the determination of the receptor-ligand interactions between IFITM proteins and SARS-CoV-2.

## INTRODUCTION

Coronaviruses are crown-shaped, single strand, positive polarity, and enveloped RNA viruses. The virus that causes COVID-19 is classified as beta coronavirus and belongs to the same subgroup as SARS. Therefore it is called SARS-CoV-2 by International Virus Taxonomy Committee [1–3]. SARS-CoV-2 interacts with angiotensin converter receptor-2 (ACE2) and binds to the cell. It uses Spike protein to binds the human lung epithelial cells. Spike protein has three chains that are A, B, and C; respectively. Spike protein plays a key role in SARS-CoV-2 infection in human cells with these three chains. It is a major antigene for attachment of the virus via ACE2 receptor to the cell [4].

Spike protein consists of two subunits. S1 subunit is responsible for host cell receptor binding and the S2 subunit is responsible for the fusion of host cell membranes. S1 domain binds the receptors and it functions as the receptor-binding domain (RBD) [5, 6]. Spike protein RBD provides the Coronaviruses binding to the host human receptors. Two types of Spike protein have been identified. The structures of the SARS-CoV-2 Spike protein were determined by cryogenic electron microscopy and it finds out two different (open and closed) conformations of the receptor-binding site. The open form is called 6VYB and the closed-form is called 6VXX. The SARS-CoV-2 Spike protein must be in the open conformation to interact with ACE2 to initiate viral entry. This open type triggers severe infection [7, 8].

Polymorphisms are genetic differences that occur with a frequency greater than 1% in a population [9]. Polymorphisms are not the cause of diseases but may be a cause of susceptibility to disease. The most common type of polymorphism in the human genome is single nucleotide polymorphisms. SNP is the difference of a single nucleotide in the DNA sequence. SNPs have a major effect on disease susceptibility and drug response. If there is a mutation such as SNPs, structural variations, chromosome number, and structural anomalies in an individual; these mutations or anomalies will change protein structure and function [10]. As a result, many diseases occur. In addition, changes in protein structure also affect the response to drugs [11]. Single nucleotide polymorphisms occurring in the structure of the Spike protein cause changes in the structure of the protein. In this case, small changes in conformation and sequence can lead to differences in the affinity of Spike proteins for ACE2 protein [12].

Interferon-induced transmembrane (IFITM) proteins are a family of small membrane proteins with immunological and developmental roles. Five types of IFITM proteins have been identified in humans: IFITM1, IFITM2, IFITM3, IFITM5, and IFITM10 [13]. The interferon-derived transmembrane protein family (IFITM/MIL/Fragilis) genes encode cell-surface proteins that can regulate cell adhesion and influence cell differentiation. IFITM1 and IFITM3 proteins expressed in primordial germ cells (PGCs) are involved in germ cell development, but their specific functions are unclear [14]. IFITM1 and IFITM3 proteins expressed in PGC are involved in germ cell development, but their specific functions are unclear [15]. IFITM1, IFITM2, and IFITM3 proteins are natal immune responses to virus infections. They regulate the fusion of the virus and direct it to lysosomes [16, 17]. IFITM3 can affect membrane stiffness and warp to inhibit virus membrane fusion [18]. This action prevents the release of viral particles into the cytoplasm [19].

IFITM proteins are a cofactor for SARS-CoV-2 infection and these are potential targets for therapeutic approaches. Peptides and targeting antibodies that derived from IFITM (especially IFITM1, 2, and 3) have important roles in inhibition of SARS-CoV-2 entry and replication in the few cell types as the host human lung cells, cardiomyocytes, and gut [20]. In this study, we demonstrated the interactions (binding energies of the protein-protein complexes and bond types) of wild and mutant types of IFITM1, IFITM2, and IFITM3 proteins with open and closed forms of SARS-CoV-2 Spike protein by molecular docking approach. The workflow chart of the study can be seen in Figure 1.

**Figure 1.**
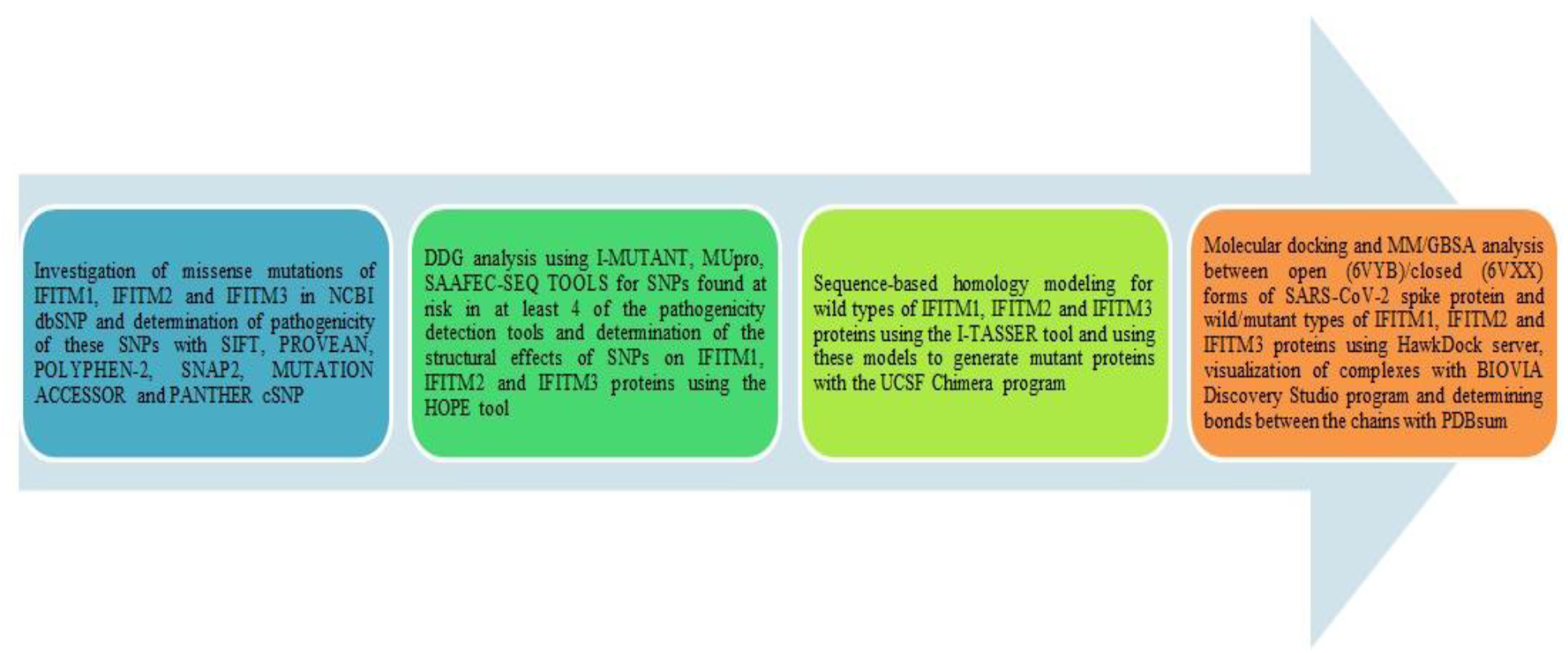
Workflow chart of the study.

## MATERIALS AND METHODS

### Data extraction and evaluation of the missense SNPs of *IFITM1, IFITM2,* and *IFITM3*

NCBI dbSNP (https://www.ncbi.nlm.nih.gov/snp/) [21] was utilized for the determination of the missense SNPs on *IFITM1*, *IFITM2*, and *IFITM3* on July 2021. The web-based tools; Sorting Intolerant From Tolerant (SIFT) (https://sift.bii.a-star.edu.sg/www/SIFT_dbSNP.html) [22], Protein Variation Effect Analyzer (PROVEAN) (http://provean.jcvi.org/seq_submit.php) [23], PolyPhen-2 (http://genetics.bwh.harvard.edu/pph2/) [24], SNAP2 (https://rostlab.org/services/snap2web/) [25], Mutation Assessor (http://mutationassessor.org/r3/) [26, 27], and Protein Analysis Through Evolutionary Relationships (PANTHER) coding SNP (cSNP) (http://www.pantherdb.org/tools/csnpScoreForm.jsp) [28] tools were used for the first analysis to determine the effects of the missense SNPs. SIFT is a program that predicts the functional effect of an amino acid substitution in a particular protein. If the score provided by the program is <= 0.05, the effect is deleterious; the exact opposite, the score is > 0.05 then the effect is determined as tolerated [29]. The rsIDs of missense SNPs extracted from NCBI dbSNP were pasted into the SIFT4G predictions tool. Besides the effects of amino acid substitutions, SIFT also provides the information of nucleotide changes and amino acid changes as a table caused by missense SNPs. PROVEAN is a web-based tool in which functional effects are determined based on the amino acid sequence of a protein found in any organism. The default score threshold of the PROVEAN Protein tool is -2.5. In the binary classification, if the score is <= -2.5, the effect is deleterious; the score is > -2.5, the effect is neutral [30]. Amino acid FASTA sequences of IFITM1, IFITM2, and IFITM3 proteins were copied from UniProt (https://www.uniprot.org/) [31] and inserted into the PROVEAN Protein tool with substitutions. PolyPhen-2 works similarly to PROVEAN. PolyPhen-2 scores differ from 0.0 (benign) to 1.0 (damaging) [32]. SNAP2 is a classifier based on a neural network and utilizes evolutionary information from multiple sequence alignments, predicted structural properties, etc. of the amino acid sequences [33]. Mutation Assessor and PANTHER cSNP tools estimate the effects of the substitutions in the evolutionary base. In the first analysis, if the prediction is possibly damaging (POD), probably damaging (PRD), deleterious (DLT), and/or the functional impacts are low, medium, or high in at least 4 of 6 tools, they were saved for the further analysis. Neutral or benign effects were not considered in the study (Table 1).

**Table 1.** Evaluation of the missense SNPs of *IFITM1*, *IFITM2*, and *IFITM3* via web-based prediction tools

| GENE | SNP | ALLELE CHANGE | AMINO ACID CHANGE | SIFT |  | PROVEAN |  | PolyPhen-2 |  | SNAP2 |  | Mutation Assessor |  | PANTHER cSNP |  |
| --- | --- | --- | --- | --- | --- | --- | --- | --- | --- | --- | --- | --- | --- | --- | --- |
|  |  |  |  | Sc | Pr | Sc | Pr | Sc | Pr | Sc | Pr | FI Sc | FUNC. IMPACT | Preservation Time (million year) | Pr |
| IFITM1 | rs9667990 | C>G | P13A | 1 | TOL | 2.676 | Neutral | 0.000 | BENIGN | EFFECT | 29 | -1.72 | neutral | 30 | PRB |
| IFITM1 | rs11552361 | A>G | N26S | 0.101 | TOL | -3.129 | DLT | 0.218 | BENIGN | EFFECT | 10 | 1.21 | low | 30 | PRB |
| IFITM1 | rs11552362 | C>G | L45V | 0.392 | TOL | -1.548 | Neutral | 0.116 | BENIGN | NEUTRAL | -49 | 1.715 | low | 91 | PRB |
| IFITM1 | rs11552366 | T>C | I87T | 0.026 | DLT | -4.579 | DLT | 0.921 | POD | EFFECT | 51 | 3.58 | high | 456 | PRD |
| IFITM1 | rs191154799 | G>A | M115I | 1 | TOL | 0.428 | Neutral | 0.001 | BENIGN | NEUTRAL | -10 | -0.345 | neutral | 30 | PRB |
| IFITM1 | rs199539158 | A>G | H113R | 0.349 | TOL | -0.401 | Neutral | 0.218 | BENIGN | EFFECT | 14 | 0.55 | neutral | 30 | PRB |
| IFITM1 | rs200518942 | G>C | Q120H | 0.017 | DLT | -1.700 | Neutral | 0.855 | POD | EFFECT | 20 | 2.085 | medium | 30 | PRB |
| IFITM1 | rs200528039 | G>A | G74R | 0.077 | TOL | -7.210 | DLT | 0.977 | PRD | EFFECT | 69 | 1.455 | low | 456 | PRD |
| IFITM1 | rs201082701 | G>A | V105I | 0.879 | TOL | -0.325 | Neutral | 0.005 | BENIGN | NEUTRAL | -10 | 0.875 | low | 31 | PRB |
| IFITM1 | rs201402251 | A>G | H28R | 0.444 | TOL | 0.369 | Neutral | 0.000 | BENIGN | NEUTRAL | -35 | 0.43 | neutral | 30 | PRB |
| IFITM1 | rs367880644 | G>A | E30K | 0.095 | TOL | -3.412 | DLT | 0.126 | BENIGN | EFFECT | 50 | 1.645 | low | 324 | POD |
| IFITM1 | rs367961349 | C>T | H6Y | 0.293 | TOL | -3.213 | DLT | 0.009 | BENIGN | EFFECT | 5 | 1.33 | low | 324 | POD |
| IFITM1 | rs368479717 | A>G | I98V | 0.312 | TOL | -0.605 | Neutral | 0.247 | BENIGN | NEUTRAL | -37 | 2.01 | medium | 324 | POD |
| IFITM1 | rs371803538 | G>A | V33M | 0.159 | TOL | -1.889 | Neutral | 0.884 | POD | NEUTRAL | -16 | 1.465 | low | 324 | POD |
| IFITM1 | rs372454990 | G>A | G107S | 0.252 | TOL | -0.321 | Neutral | 0.089 | BENIGN | EFFECT | 18 | 1.82 | low | 30 | PRB |
| IFITM1 | rs373112031 | G>A | V61M | 0.11 | TOL | -1.932 | Neutral | 0.956 | POD | EFFECT | 10 | 1.795 | low | 324 | POD |
| IFITM1 | rs373832568 | G>A | V72M | 0.403 | TOL | -0.066 | Neutral | 0.061 | BENIGN | EFFECT | 6 | -0.28 | neutral | 31 | PRB |
| IFITM1 | rs374294080 | G>A | V24M | 0.174 | TOL | -2.081 | Neutral | 0.631 | POD | EFFECT | 29 | 1.245 | low | 324 | POD |
| IFITM1 | rs374941646 | G>A | C84Y | 0.14 | TOL | -9.698 | DLT | 0.952 | POD | EFFECT | 47 | 3.265 | medium | 324 | POD |
| IFITM1 | rs377130310 | T>C | L41P | 0.024 | DLT | -6.476 | DLT | 0.574 | POD | EFFECT | 72 | 3.485 | medium | 324 | POD |
| IFITM1 | rs377178214 | C>T | P34S | 0.016 | DLT | -6.093 | DLT | 0.998 | PRD | EFFECT | 60 | 3.565 | high | 324 | POD |
| IFITM2 | rs14408 | T>C | M41T | 1 | TOL | 3.282 | Neutral | 0.000 | BENIGN | EFFECT | 24 | -1.61 | neutral | 1 | PRB |
| IFITM2 | rs14408 | T>C | M21T | 1 | TOL | -3.445 | DLT | 0.006 | BENIGN | EFFECT | 19 | 0.985 | low | 1 | PRB |
| IFITM2 | rs1058900 | T>C | V33A | 1 | TOL | 1.728 | Neutral | 0.000 | BENIGN | EFFECT | 24 | -0.48 | neutral | 1 | PRB |
| IFITM2 | rs1059091 | A>C | I121L | 0.103 | TOL | -1.002 | Neutral | 0.000 | BENIGN | EFFECT | 85 | 3.185 | medium | 1 | PRB |
| IFITM2 | rs1059091 | A>G | I121V | 0.13 | TOL | -0.322 | Neutral | 0.000 | BENIGN | NEUTRAL | -33 | 2.215 | medium | 1 | PRB |
| IFITM2 | rs74901761 | T>C | F115L | 1 | TOL | 1.008 | Neutral | 0.000 | BENIGN | NEUTRAL | -28 | -0.46 | neutral | 1 | PRB |
| IFITM2 | rs75054971 | G>A | V53M | 0.15 | TOL | -2.272 | Neutral | 0.766 | POD | EFFECT | 13 | 1.415 | low | 325 | POD |
| IFITM2 | rs75054971 | G>A | V33M | 0.161 | TOL | -1.078 | Neutral | 0.574 | POD | EFFECT | 76 | 1.545 | low | 1 | PRB |
| IFITM2 | rs183593068 | G>A | M116I | 0.396 | TOL | -1.342 | Neutral | 0.013 | BENIGN | NEUTRAL | -2 | 1.6 | low | 1 | PRB |
| IFITM2 | rs199693024 | T>C | V5A | 0.373 | TOL | 0.039 | Neutral | 0.038 | BENIGN | EFFECT | 5 | 1.935 | low | 1 | PRB |
| IFITM2 | rs201079035 | G>A | G14S | 0.504 | TOL | -1.333 | Neutral | 0.014 | BENIGN | EFFECT | 21 | 0.955 | low | 1 | PRB |
| IFITM2 | rs202139418 | A>C | T69P | 0.25 | TOL | -0.654 | Neutral | 0.000 | BENIGN | EFFECT | 49 | -0.835 | neutral | 1 | PRB |
| IFITM2 | rs368808923 | G>A | E20K | 0.128 | TOL | -2.779 | DLT | 0.950 | POD | EFFECT | 56 | 2.5 | medium | 1 | PRB |
| IFITM2 | rs369960951 | A>G | M30V | 1 | TOL | 1.627 | Neutral | 0.000 | BENIGN | NEUTRAL | -10 | -1.82 | neutral | 1 | PRB |
| IFITM2 | rs371179996 | A>G | N63S | 0.101 | TOL | -4.219 | DLT | 0.986 | PRD | EFFECT | 16 | 2.09 | medium | 456 | PRD |
| IFITM2 | rs371346866 | G>A | E25K | 0.161 | TOL | -3.492 | DLT | 0.218 | BENIGN | EFFECT | 58 | 2.445 | medium | 1 | PRB |
| IFITM2 | rs371736596 | C>A | P17H | 0.063 | TOL | -4.344 | DLT | 0.030 | BENIGN | EFFECT | 57 | 2.3 | medium | 176 | PRB |
| IFITM2 | rs373014074 | C>G | R132G | 0.006 | DLT | -0.658 | Neutral | 0.003 | BENIGN | EFFECT | 72 | 0.69 | neutral | 1 | PRB |
| IFITM2 | rs373569657 | G>A | M30I | 0.236 | TOL | 0.744 | Neutral | 0.000 | BENIGN | EFFECT | 20 | 0 | neutral | 1 | PRB |
| IFITM2 | rs376286409 | G>A | R132Q | 0.233 | TOL | -0.275 | Neutral | 0.001 | BENIGN | EFFECT | 72 | -0.41 | neutral | 1 | PRB |
| IFITM2 | rs376647548 | G>C | Q26H | 1 | TOL | 4.875 | Neutral | 0.000 | BENIGN | NEUTRAL | -46 | -1.425 | neutral | 1 | PRB |
| IFITM2 | rs376647548 | G>C | Q6H | 1 | TOL | -1.691 | Neutral | 1.000 | PRD | EFFECT | 18 | 2.125 | medium | 1 | PRB |
| IFITM3 | rs1136853 | G>T | H3Q | 0.008 | DLT | -2.939 | DLT | 0.437 | BENIGN | EFFECT | 76 | 2.815 | medium | 97 | PRB |
| IFITM3 | rs199582787 | C>T | V31M | 0.124 | TOL | -1.677 | Neutral | 0.019 | BENIGN | EFFECT | 2 | 1.445 | low | 324 | POD |
| IFITM3 | rs199749095 | G>T | P70T | 0.391 | TOL | 0.654 | Neutral | 0.000 | BENIGN | NEUTRAL | -1 | 0.865 | low | 31 | PRB |
| IFITM3 | rs200737925 | T>A | H57L | 0.005 | DLT | -10.664 | DLT | 0.759 | POD | EFFECT | 77 | 3.75 | high | 324 | POD |
| IFITM3 | rs200737925 | T>A | H36L | 0.006 | DLT | -7.090 | DLT | 0.097 | BENIGN | EFFECT | 50 | 3.065 | medium | 97 | PRB |
| IFITM3 | rs368803166 | C>G | S50T | 0.273 | TOL | -2.106 | Neutral | 0.931 | POD | EFFECT | 17 | 2.035 | medium | 324 | POD |
| IFITM3 | rs370158790 | G>A | A34V | 0.326 | TOL | -1.828 | Neutral | 0.005 | BENIGN | EFFECT | 10 | 0.4 | neutral | 324 | POD |
| IFITM3 | rs371741630 | C>T | V54M | 0.14 | TOL | -2.272 | Neutral | 0.965 | PRD | NEUTRAL | -20 | 2 | medium | 324 | POD |
| IFITM3 | rs376417425 | C>T | V90I | 0.138 | TOL | -0.912 | Neutral | 1.000 | PRD | EFFECT | 14 | 2.645 | medium | 324 | POD |

Sc: Score, Pr: Prediction, TOL: Tolerated, DLT: Deleterious, POD: Possibly Damaging, PRD: Probably Damaging.

### DDG determination and structural Analysis of the pathogenic forms of IFITM1, IFITM2, and IFITM3 proteins

DDG (or referred to as ΔΔG, double changes in Gibbs free energy)s and stability changes of the proteins were determined in the web-based prediction tools: I-Mutant (http://gpcr2.biocomp.unibo.it/cgi/predictors/I-Mutant3.0/I-Mutant3.0.cgi) [34], MUpro (http://mupro.proteomics.ics.uci.edu/) [35] and SAAFEC-SEQ (http://compbio.clemson.edu/SAAFEC-SEQ/#started) [36]. Default settings (Temperature: 25 ^0^C, pH: 7.0) and the “DDG Value and Binary Classification” option were preferred in I-Mutant. Sequence-based analyses were performed for all pathogenic proteins in the aforementioned tools. If two or three of I-Mutant, MUpro, and SAAFEC-SEQ give the same result, the mutation is considered to make the protein more stable or unstable (Table 2). Also, the HOPE tool was used for structural analysis in terms of size, charge, and physical state of the proteins [37] (Table 3).

**Table 2.** DDG determination of the pathogenic forms of IFITM1, IFITM2, and IFITM3 proteins

| GENE | SNP ID | ALLELE CHANGE | AMINO ACID CHANGE | I-Mutant |  | MUpro |  | SAAFEC-SEQ |  |
| --- | --- | --- | --- | --- | --- | --- | --- | --- | --- |
|  |  |  |  | DDG (kcal/mol) | Prediction | DDG (kcal/mol) | Prediction | ddG (unit) | Prediction |
| IFITM1 | rs11552366 | T>C | I87T | -1.91 | Decrease | -2.0309983 | Decrease | -1.45 | Destabilizing |
| IFITM1 | rs200518942 | G>C | Q120H | -0.61 | Decrease | -0.59799326 | Decrease | -0.04 | Destabilizing |
| IFITM1 | rs200528039 | G>A | G74R | -0.47 | Decrease | -0.1322157 | Decrease | -0.93 | Destabilizing |
| IFITM1 | rs367880644 | G>A | E30K | -0.80 | Decrease | -1.0887728 | Decrease | -0.09 | Destabilizing |
| IFITM1 | rs367961349 | C>T | H6Y | 0.44 | Increase | -0.17753438 | Decrease | -0.59 | Destabilizing |
| IFITM1 | rs373112031 | G>A | V61M | -1.36 | Decrease | -0.70976867 | Decrease | -0.58 | Destabilizing |
| IFITM1 | rs374294080 | G>A | V24M | -1.11 | Decrease | -0.89921065 | Decrease | -0.82 | Destabilizing |
| IFITM1 | rs374941646 | G>A | C84Y | 0.02 | Increase | -0.65558741 | Decrease | -1.01 | Destabilizing |
| IFITM1 | rs377130310 | T>C | L41P | -1.54 | Decrease | -2.4357497 | Decrease | -1.10 | Destabilizing |
| IFITM1 | rs377178214 | C>T | P34S | -1.67 | Decrease | -1.3598958 | Decrease | -0.60 | Destabilizing |
| IFITM2 | rs75054971 | G>A | V53M | -1.13 | Decrease | -1.07739 | Decrease | -0.36 | Destabilizing |
| IFITM2 | rs368808923 | G>A | E20K | -0.70 | Decrease | -0.98470021 | Decrease | -0.76 | Destabilizing |
| IFITM2 | rs371179996 | A>G | N63S | -0.48 | Decrease | -1.0517659 | Decrease | -0.26 | Destabilizing |
| IFITM3 | rs1136853 | G>T | H3Q | -0.19 | Decrease | -0.51910301 | Decrease | -0.15 | Destabilizing |
| IFITM3 | rs200737925 | T>A | H57L | 0.31 | Increase | 0.1001401 | Increase | -0.10 | Destabilizing |
| IFITM3 | rs200737925 | T>A | H36L | 0.62 | Increase | 0.69450667 | Increase | -0.03 | Destabilizing |
| IFITM3 | rs368803166 | C>G | S50T | -0.39 | Decrease | -0.96036192 | Decrease | -0.07 | Destabilizing |
| IFITM3 | rs376417425 | C>T | V90I | -0.52 | Decrease | -0.25157544 | Decrease | 0.10 | Stabilizing |

**Table 3.** Structural analysis of the pathogenic forms of IFITM1, IFITM2, and IFITM3 proteins via HOPE

| GENE | SNP ID | ALLELE CHANGE | AMINO ACID CHANGE | WILD TYPE |  |  | MUTANT TYPE |  |  |
| --- | --- | --- | --- | --- | --- | --- | --- | --- | --- |
|  |  |  |  | Size | Charge | Physical State | Size | Charge | Physical State |
| IFITM1 | rs11552366 | T>C | I87T | bigger | no information | more hydrophobic | smaller | no information | less hydrophobic |
| IFITM1 | rs200518942 | G>C | Q120H | smaller | no information | no information | bigger | no information | no information |
| IFITM1 | rs200528039 | G>A | G74R | smaller | neutral | more hydrophobic | bigger | positive | less hydrophobic |
| IFITM1 | rs367880644 | G>A | E30K | smaller | negative | no information | bigger | positive | no information |
| IFITM1 | rs367961349 | C>T | H6Y | smaller | no information | less hydrophobic | bigger | no information | more hydrophobic |
| IFITM1 | rs373112031 | G>A | V61M | smaller | no information | no information | bigger | no information | no information |
| IFITM1 | rs374294080 | G>A | V24M | smaller | no information | no information | bigger | no information | no information |
| IFITM1 | rs374941646 | G>A | C84Y | smaller | no information | more hydrophobic | bigger | no information | less hydrophobic |
| IFITM1 | rs377130310 | T>C | L41P | bigger | no information | no information | smaller | no information | no information |
| IFITM1 | rs377178214 | C>T | P34S | bigger | no information | more hydrophobic | smaller | no information | less hydrophobic |
| IFITM2 | rs75054971 | G>A | V53M | smaller | no information | no information | bigger | no information | no information |
| IFITM2 | rs368808923 | G>A | E20K | smaller | negative | no information | bigger | positive | no information |
| IFITM2 | rs371179996 | A>G | N63S | bigger | no information | less hydrophobic | smaller | no information | more hydrophobic |
| IFITM3 | rs1136853 | G>T | H3Q | bigger | no information | no information | smaller | no information | no information |
| IFITM3 | rs200737925 | T>A | H57L | bigger | no information | less hydrophobic | smaller | no information | more hydrophobic |
| IFITM3 | rs200737925 | T>A | H36L | bigger | no information | less hydrophobic | smaller | no information | more hydrophobic |
| IFITM3 | rs368803166 | C>G | S50T | smaller | no information | no information | bigger | no information | no information |
| IFITM3 | rs376417425 | C>T | V90I | smaller | no information | no information | bigger | no information | no information |

### Homology modeling of wild and mutant Types of IFITM1, IFITM2, and IFITM3 proteins

Amino acid sequences (FASTA format) of wild-type protein products of *IFITM1, IFITM2,* and *IFITM3* were obtained from UniProt [31]. UniProt IDs for IFITM1, IFITM2, and IFITM3 proteins are P13164, Q01629, and Q01628, respectively. Homology modeling of these proteins was performed via the I-TASSER online server. I-TASSER provides the Confidence (C)-score, estimated TM-score, and estimated RMSD value for each model it creates. The C-score takes a value in the range of [-5,2] and a high C-score indicates high confidence for the related model [38, 39]. Also, SWISS-MODEL QMEAN (Qualitative Model Energy Analysis) [40] and ERRAT [41] scores were used for structure assessment. The quality of the model is low when the QMEAN score is less than -4 [42]. If the ERRAT score is higher than 50, the model can be accepted to be of high quality [43]. Mutant-type proteins were generated with the UCSF Chimera program [44]. Simply the wild-type residues were replaced with the mutant-type residues by entering a specific command (swapaa). Swapaa utilizes the information from a rotamer library for this operation [45].

### Molecular docking of protein-protein complexes

Molecular docking was carried out between the open and closed forms of Spike proteins of SARS-CoV-2 and wild and pathogenic mutants of the IFITM1, IFITM2, and IFITM3 proteins. For this purpose, the HawkDock server was used for the prediction and analysis of protein-protein complexes [46]. The PDB ID and chain information of the open and closed forms of the Spike proteins were entered into the server (6VYB:A for open form, 6VXX:A for closed-form) as the receptors. PDB files of wild and mutant types of IFITM1, IFITM2, and IFITM3 proteins were uploaded to the server as ligands. Hawkdock server proposes that the larger proteins be designated as receptors (http://cadd.zju.edu.cn/hawkdock/). After molecular docking is complete, Hawkdock presents 100 models for each complex and the scores of the ten of these models with the most negative scores. At this stage, the PDB files of the complexes with the most negative score were saved. MM/GBSA analyzes were performed in the same software and the binding energies were calculated for each complex. The interactions in the protein-protein complexes were visualized with the BIOVIA Discovery Studio 2021 program [47] (Figure 2, Figure 3, and Figure 4) and the H-bond option was selected for the surface display. In addition, all bonds (H-bonds, salt bridges, and non-contact bonds) and the number of these bonds in protein-protein complexes were revealed with the PDBsum online tool (https://www.ebi.ac.uk/thornton-srv/databases/pdbsum/Generate.html) [48].

**Figure 2.**
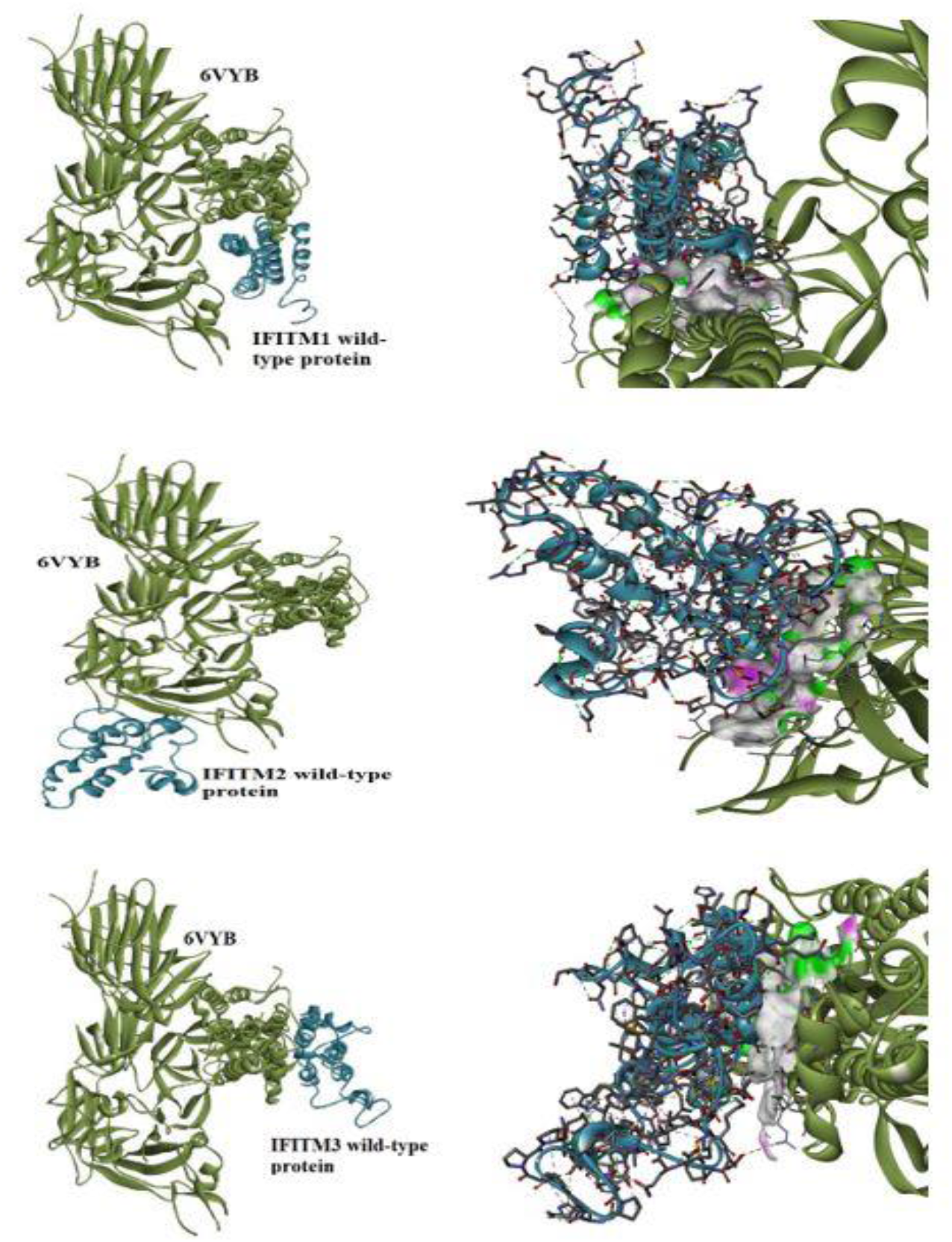
Visualization of the 6VYB-IFITM1/2/3 (wild) complexes via BIOVIA Discovery Studio.

**Figure 3.**
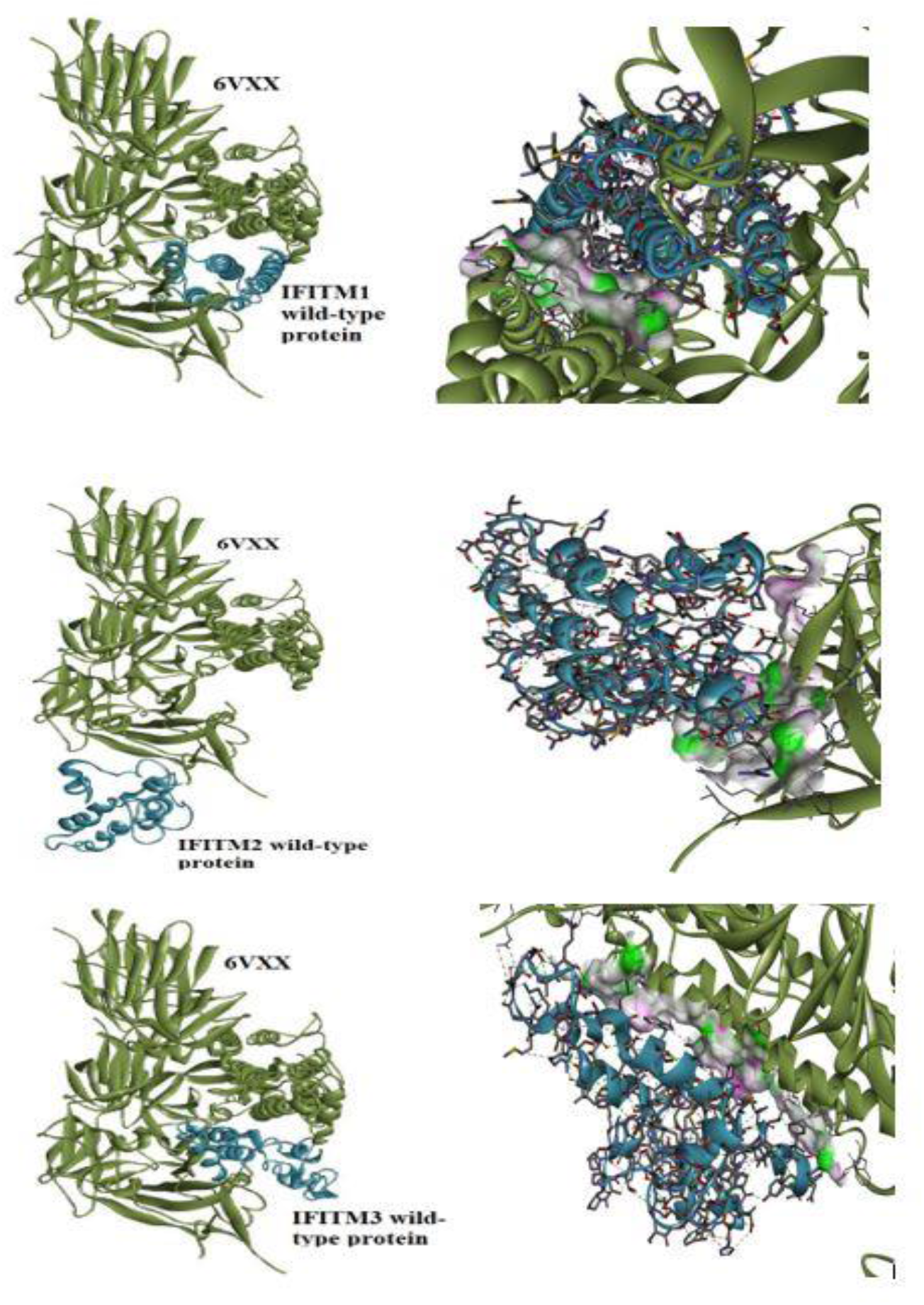
Visualization of the 6VXX-IFITM1/2/3 (wild) complexes via BIOVIA Discovery Studio.

**Figure 4.**
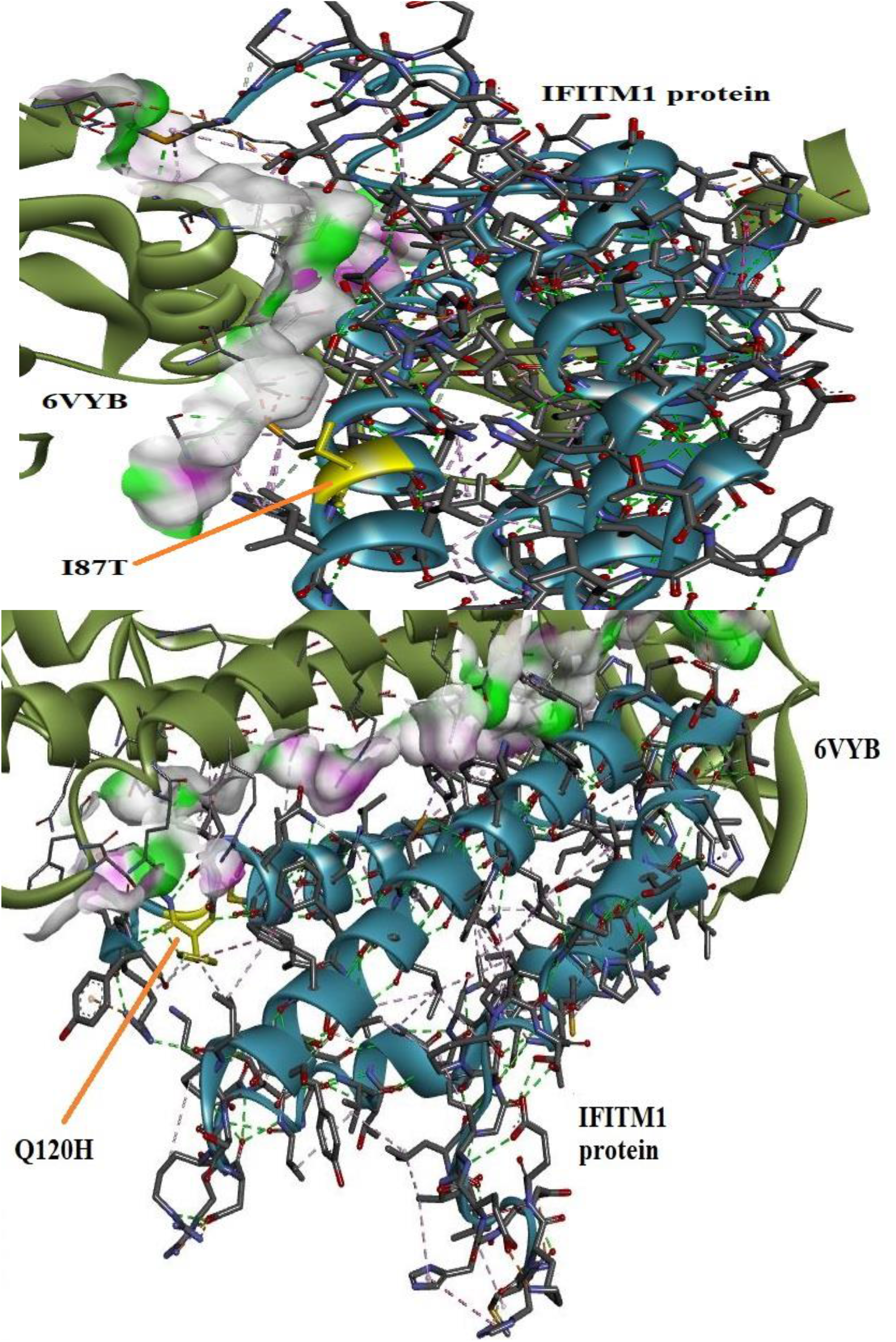
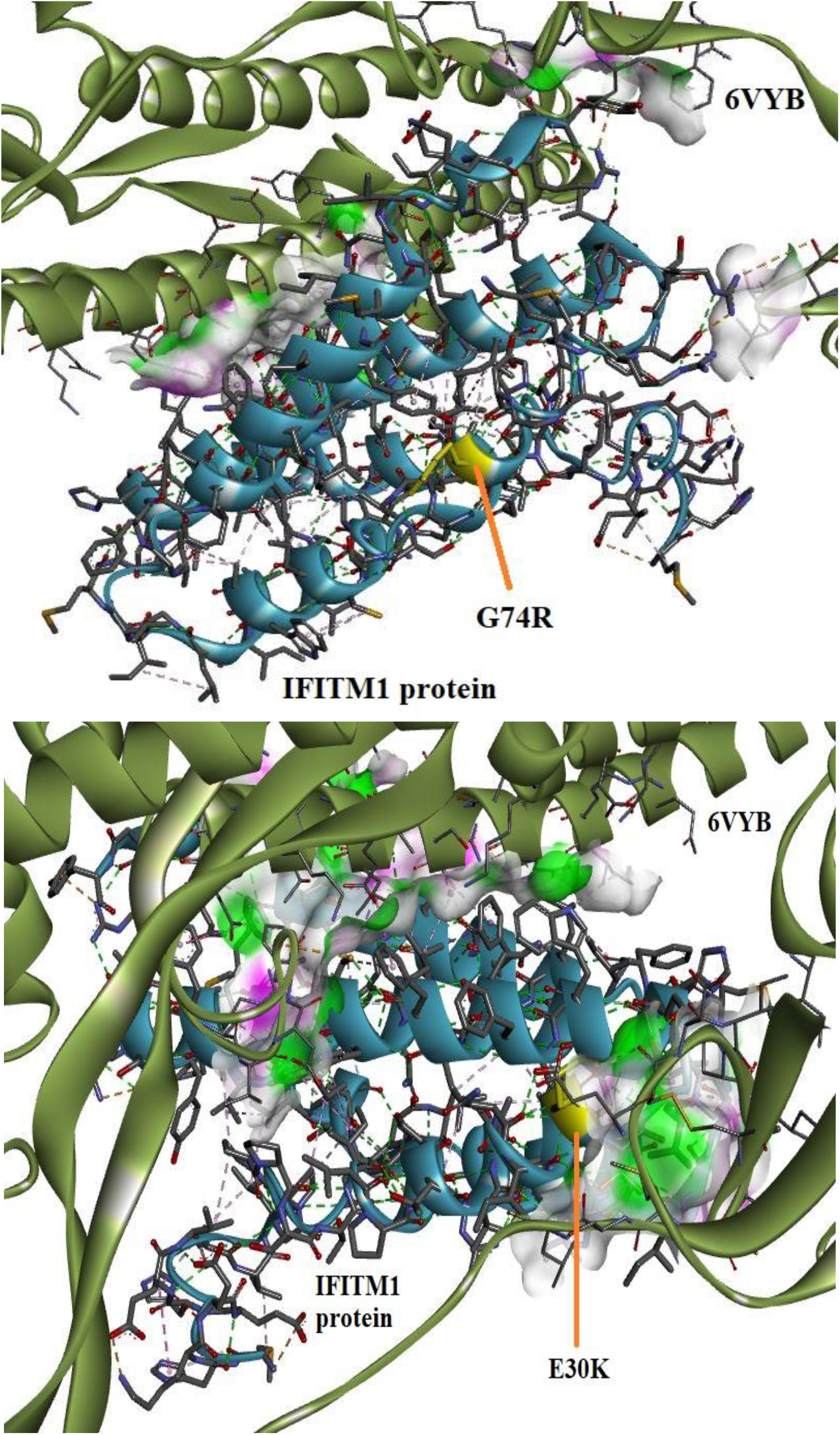
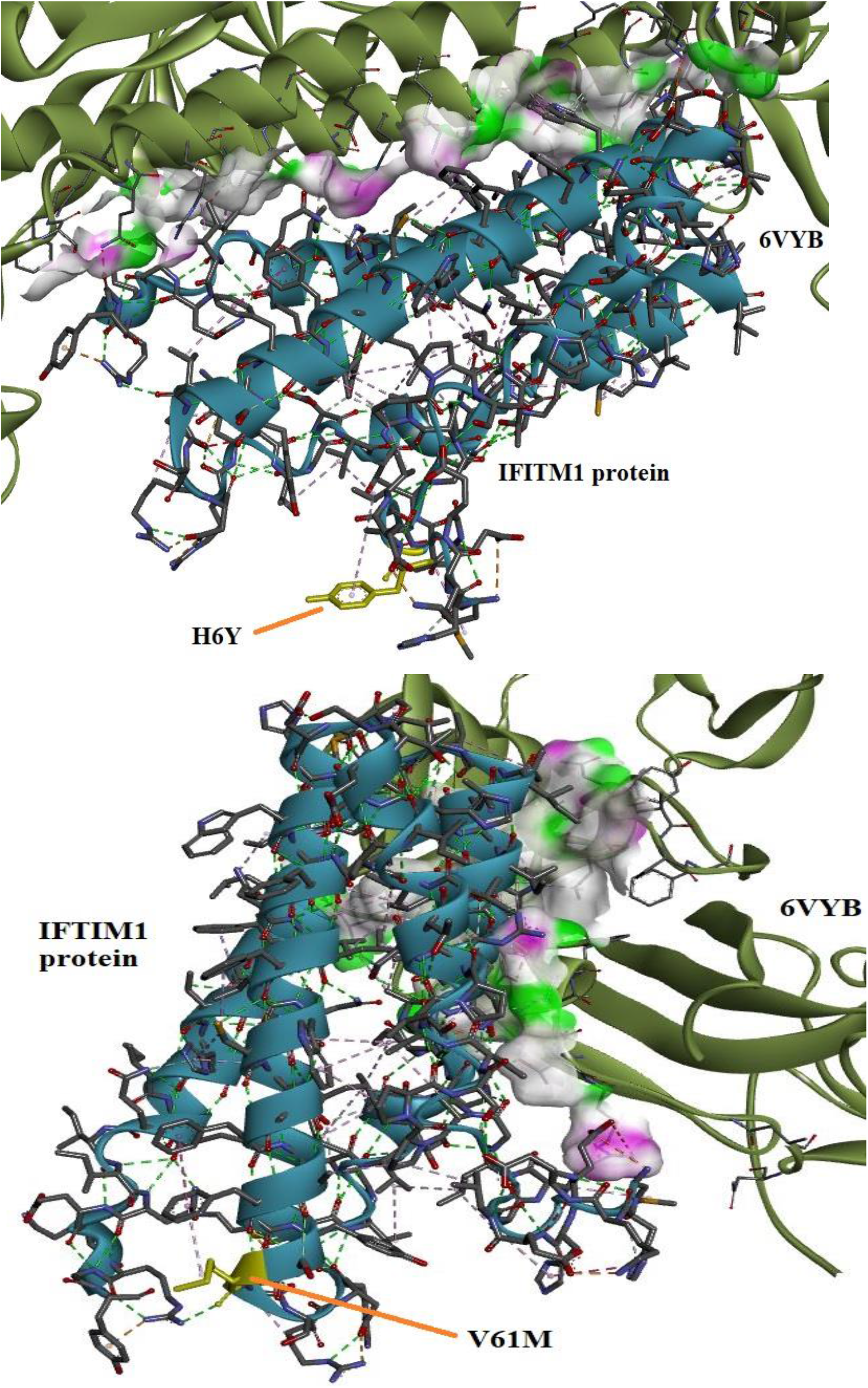
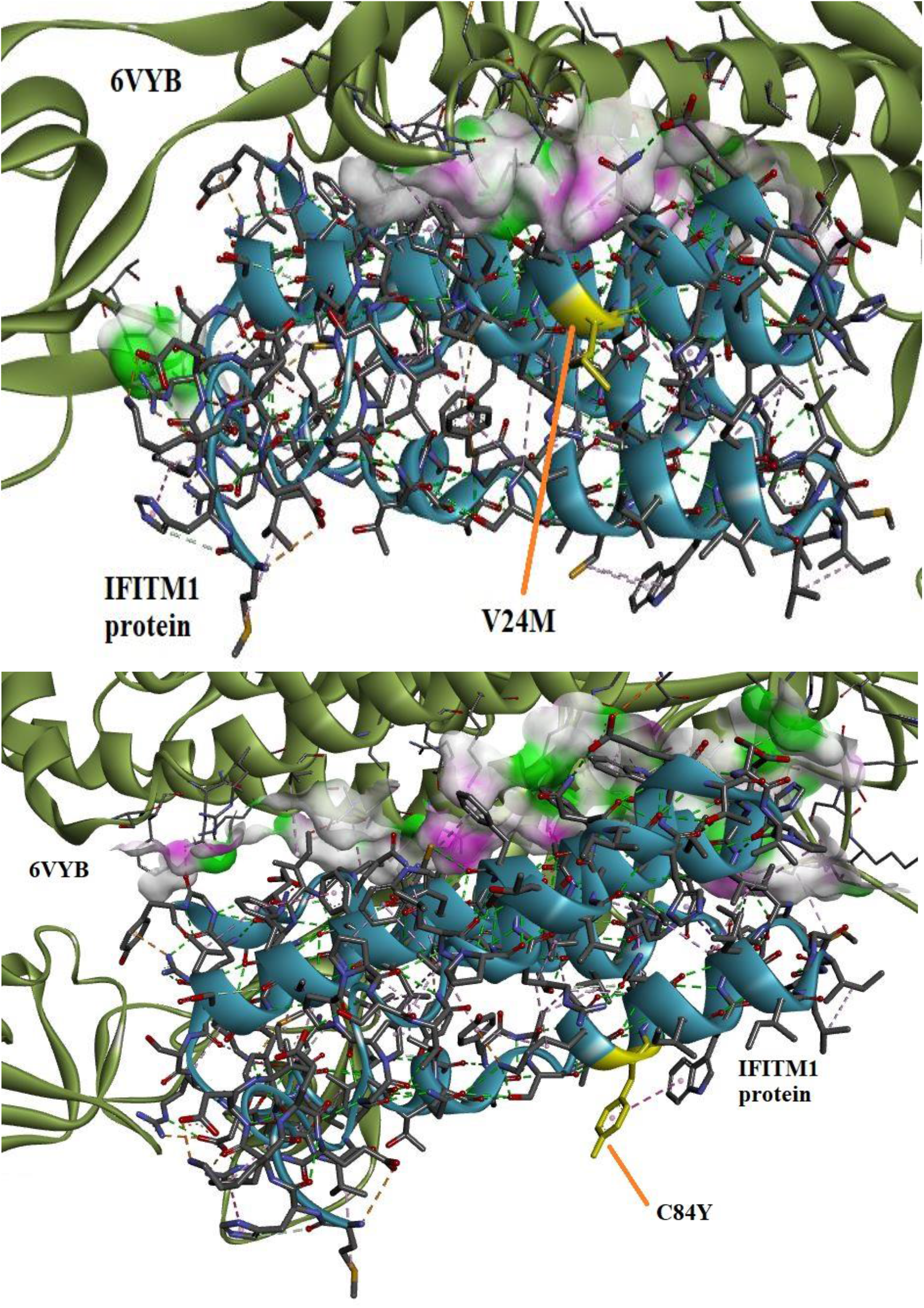
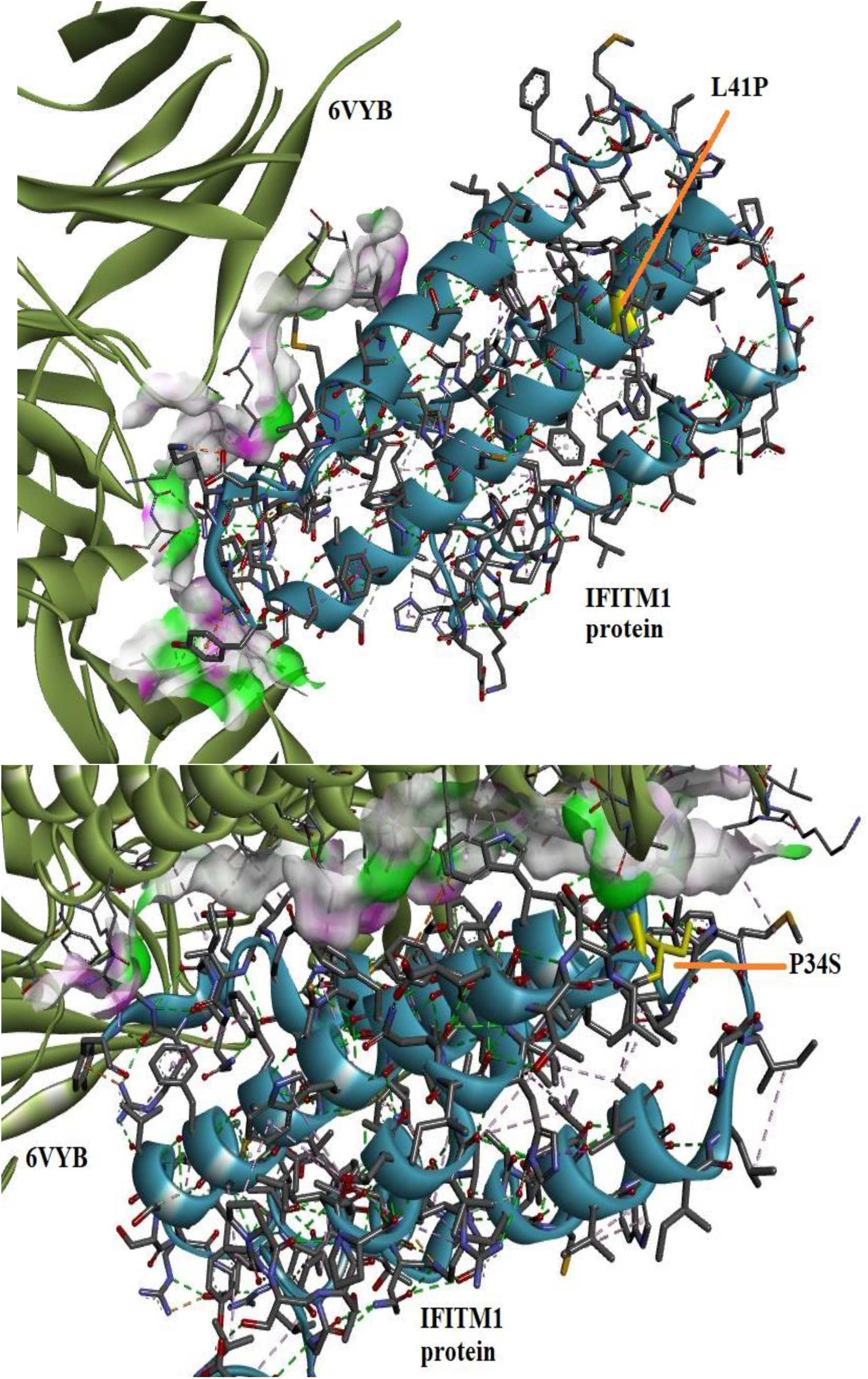
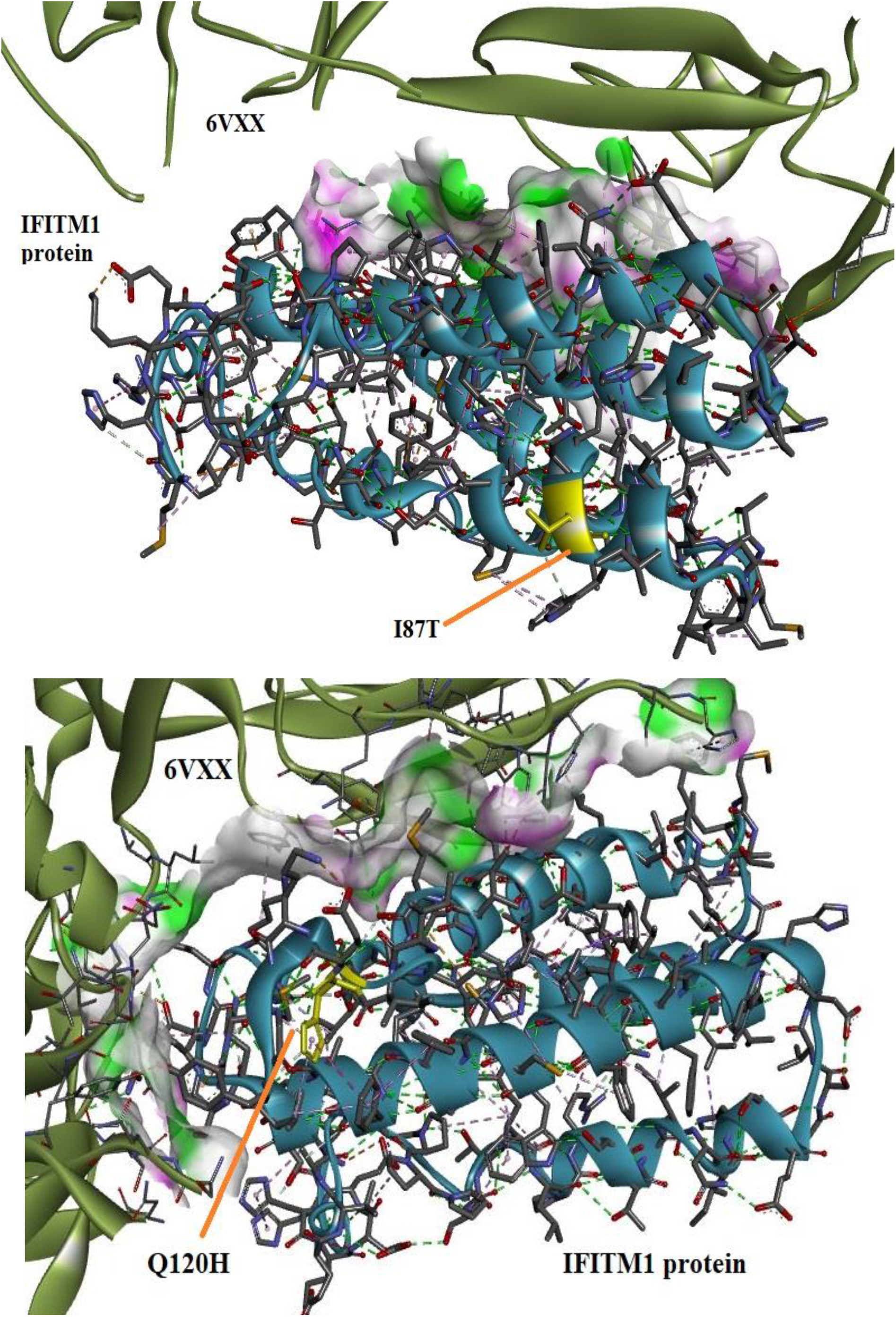
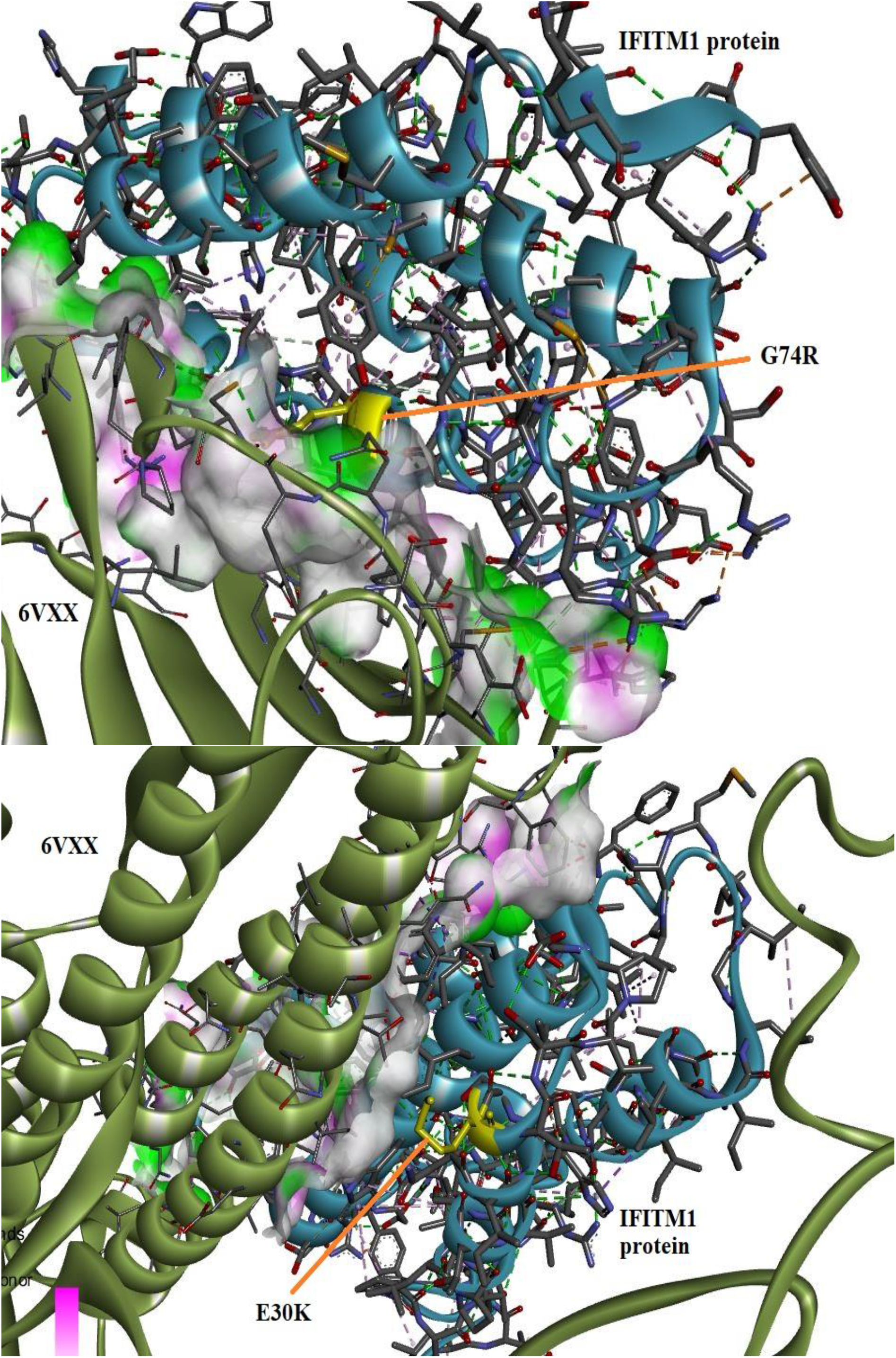
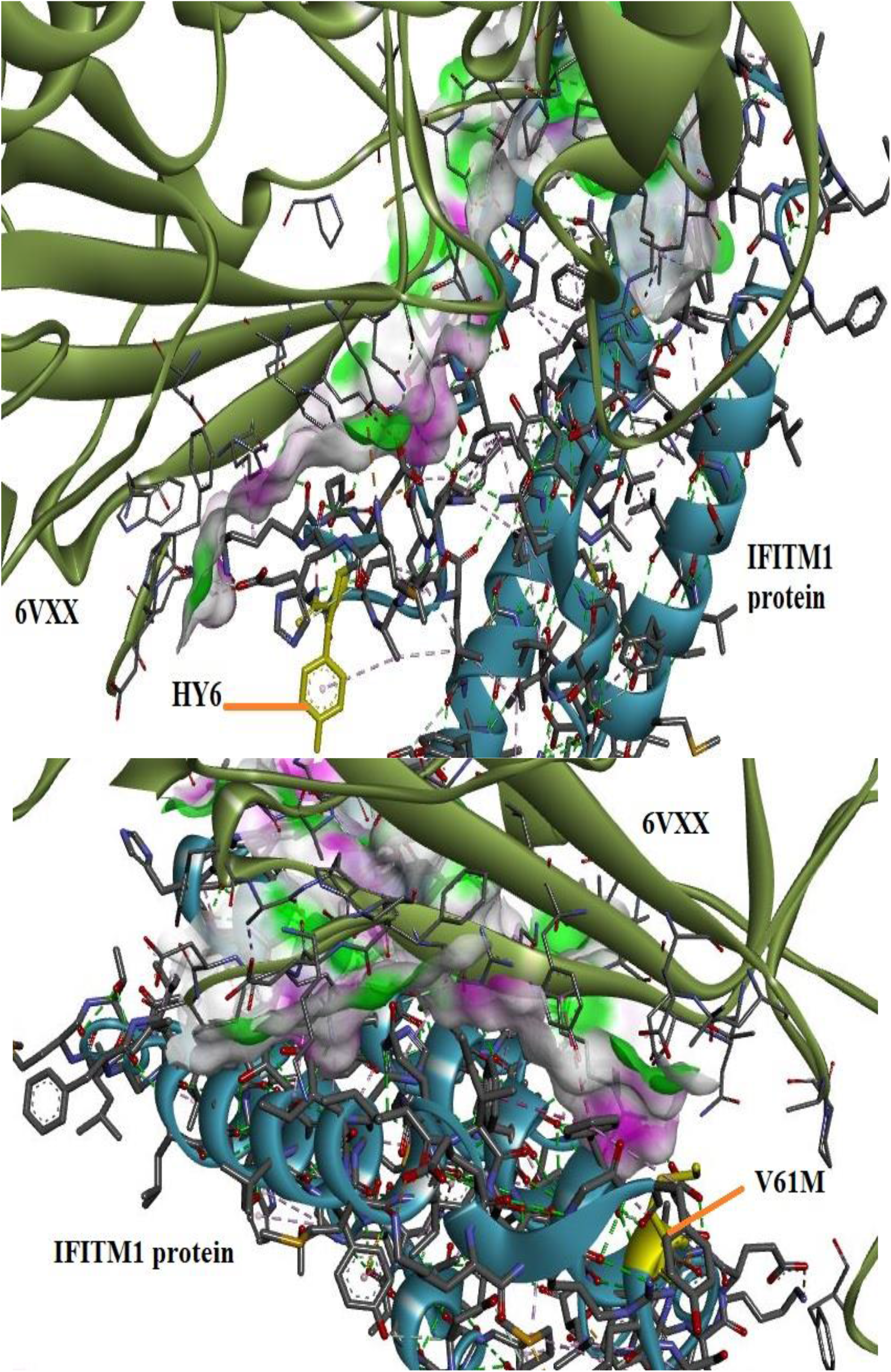
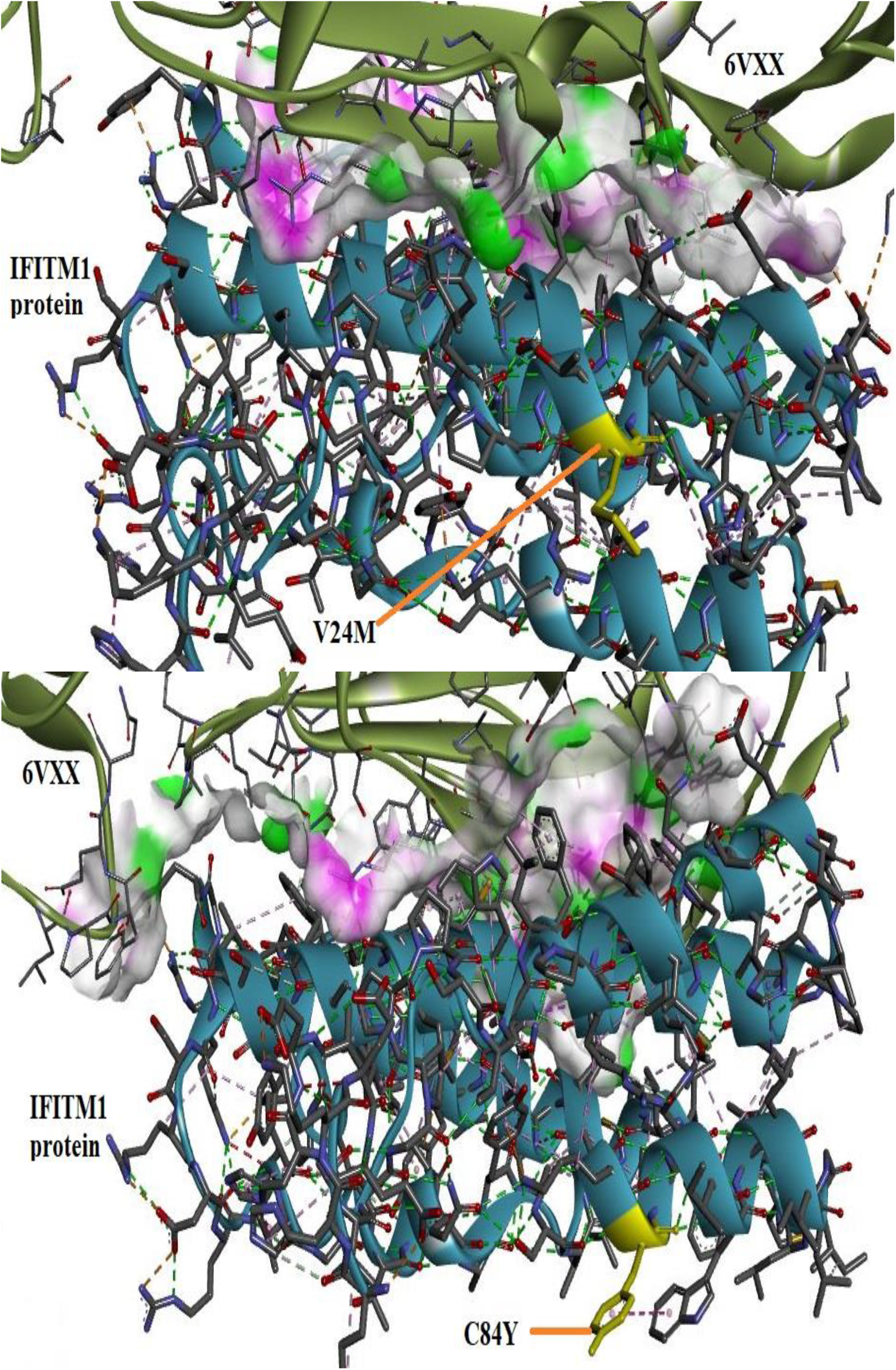
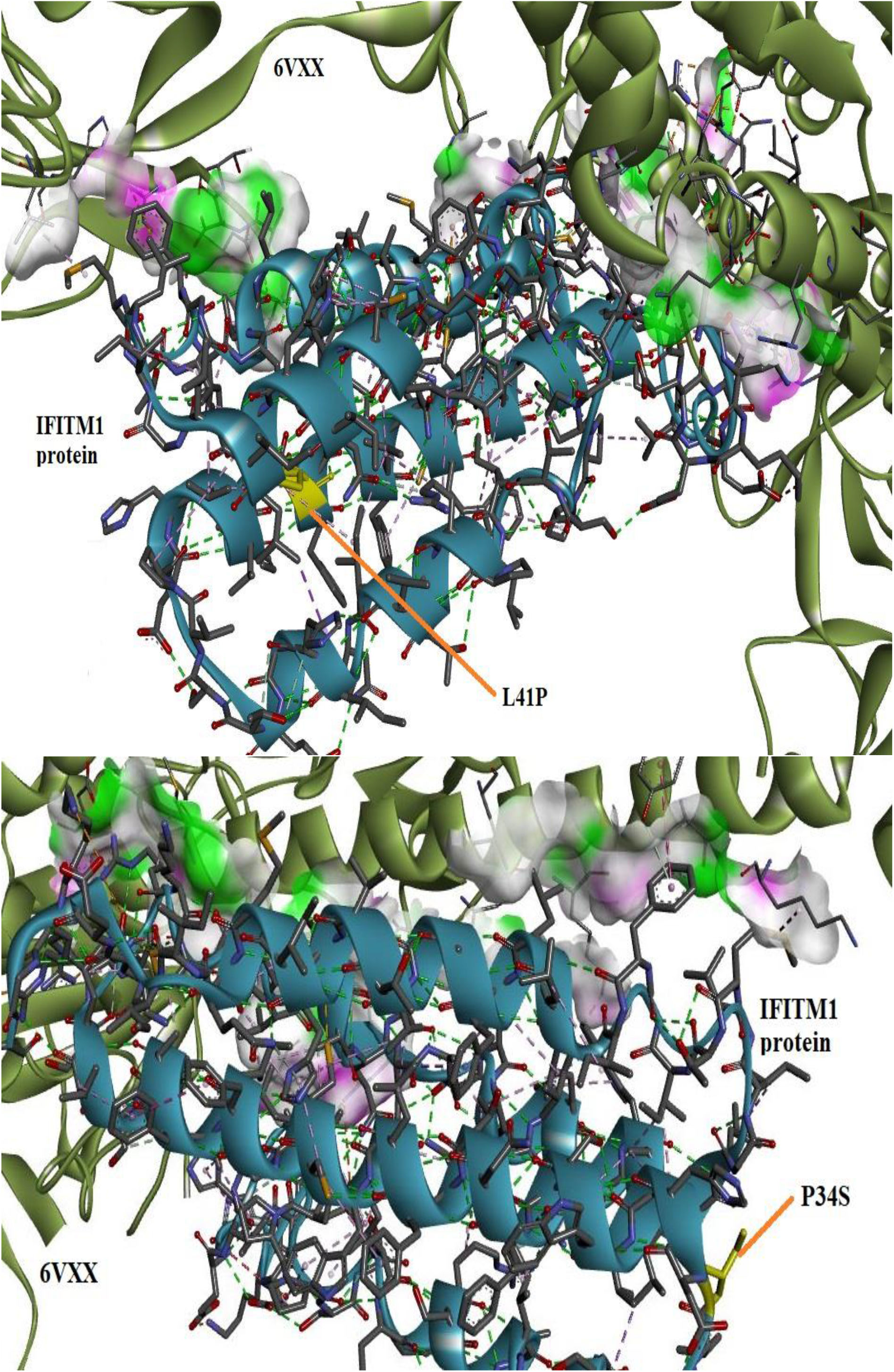
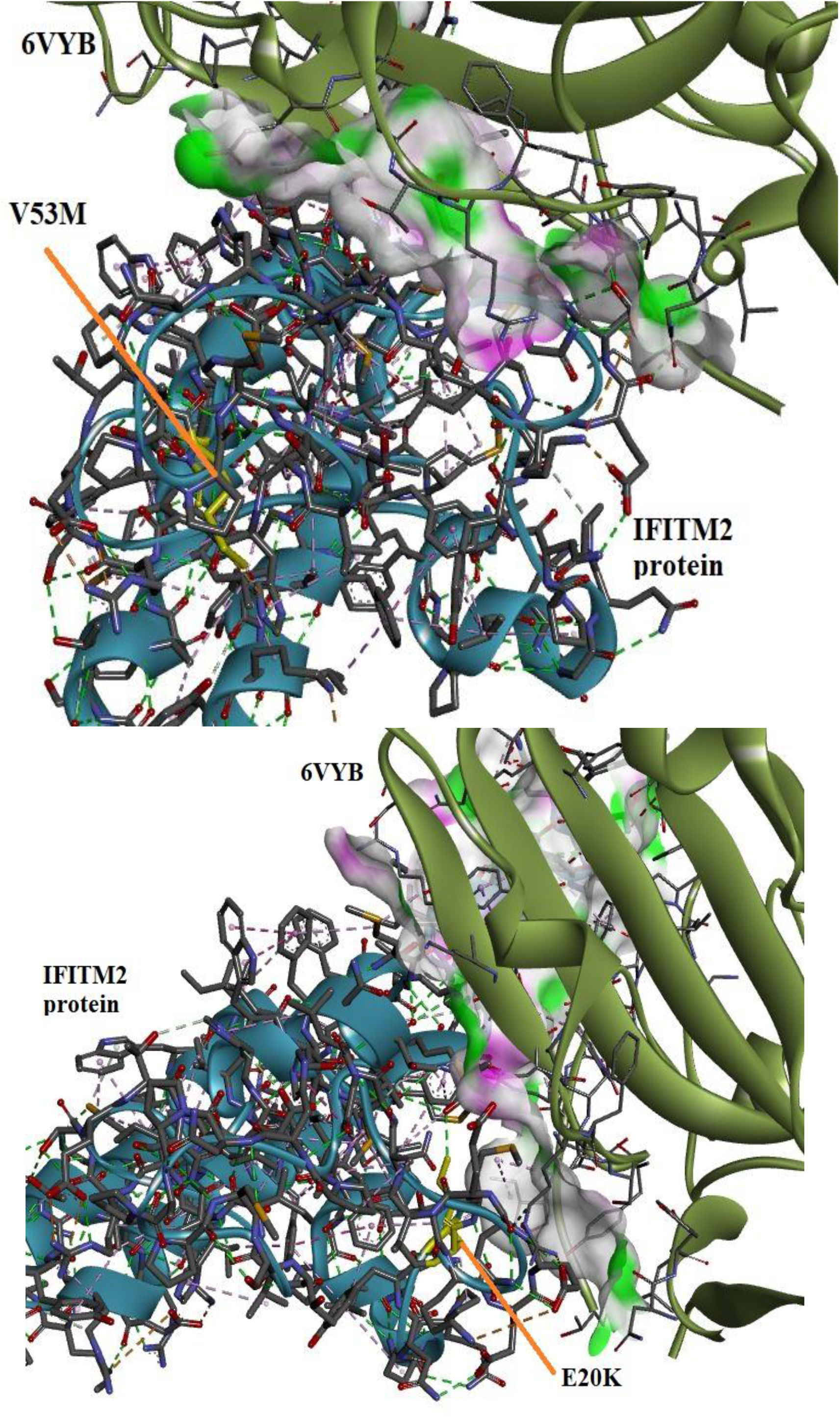
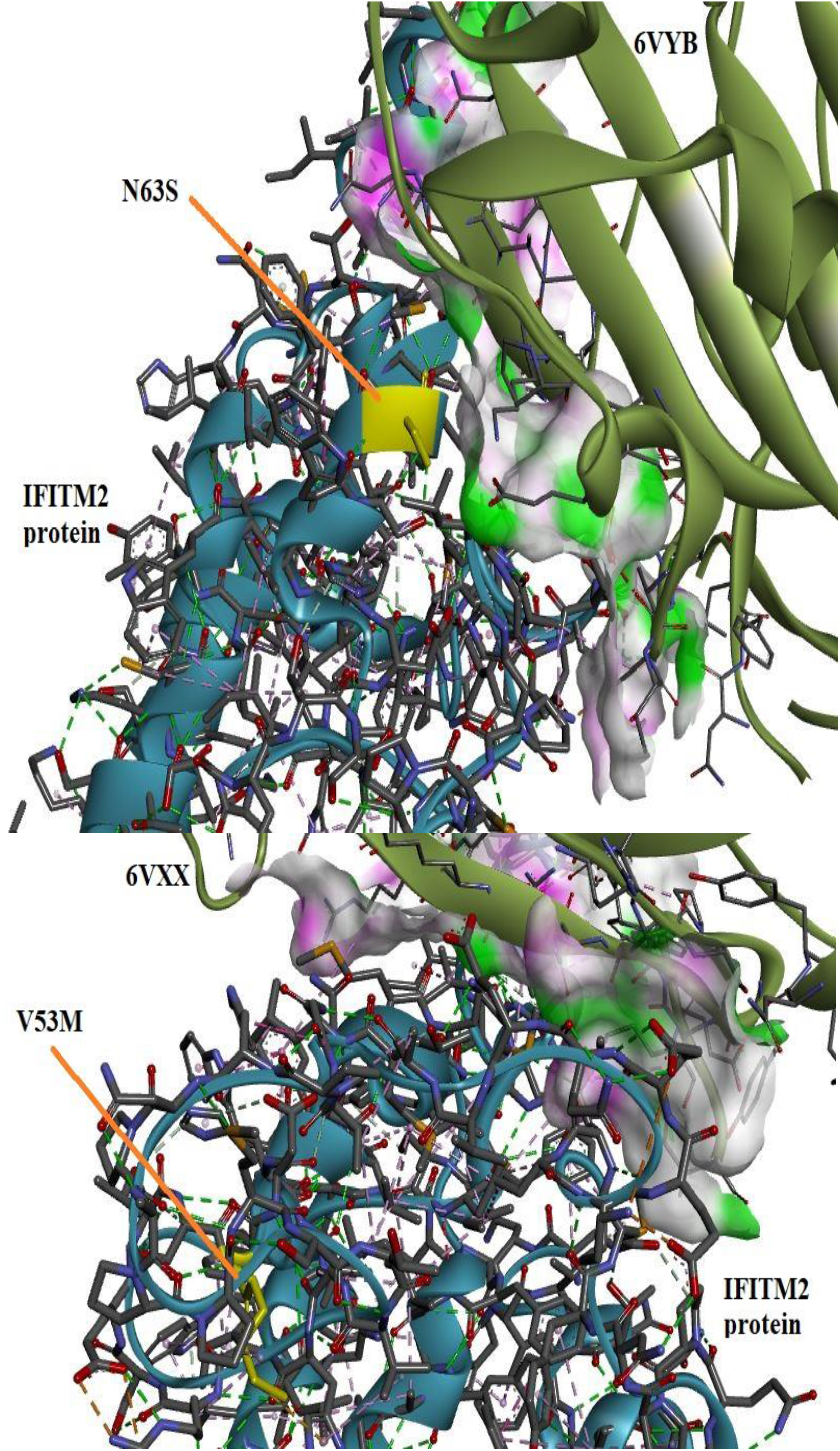
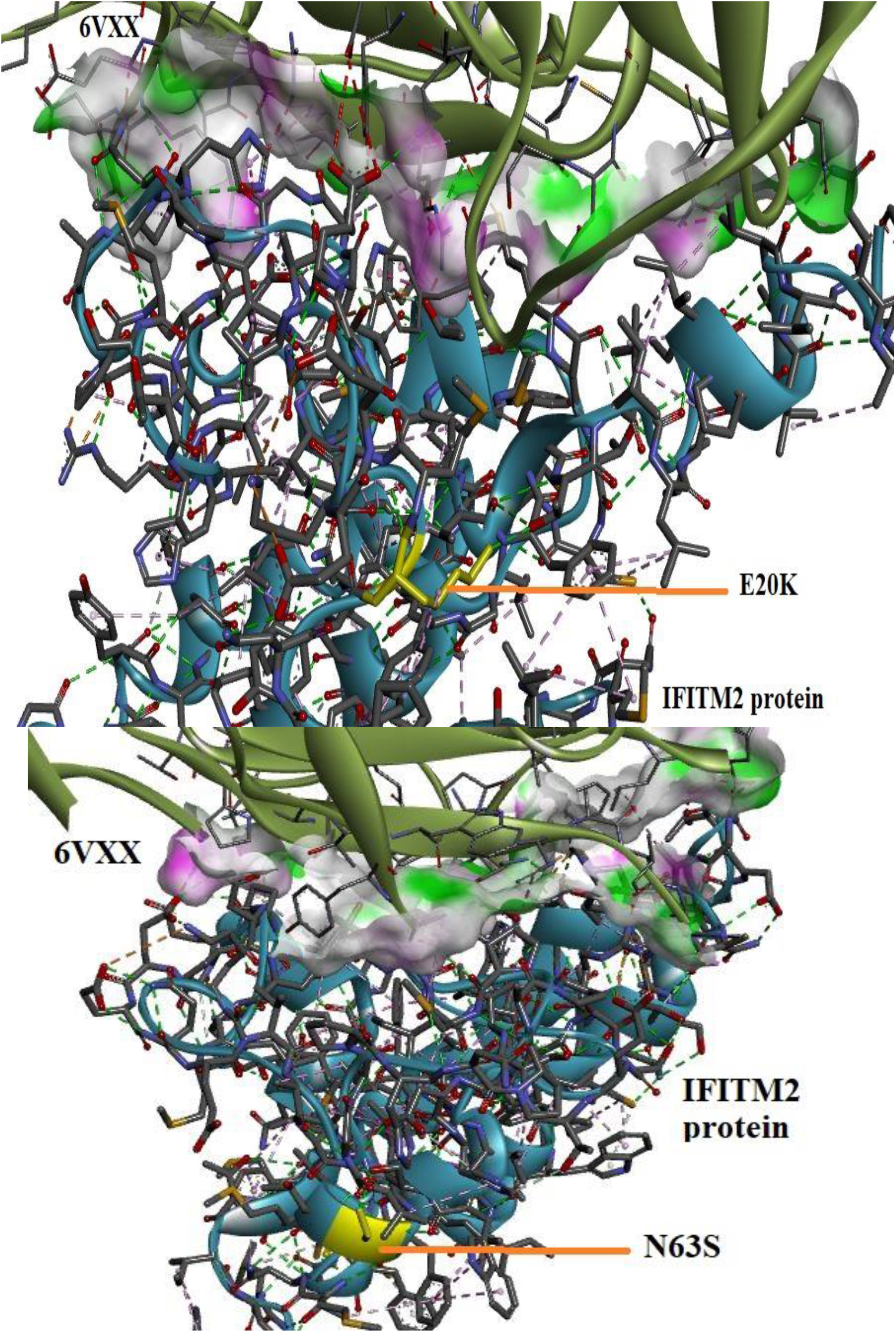
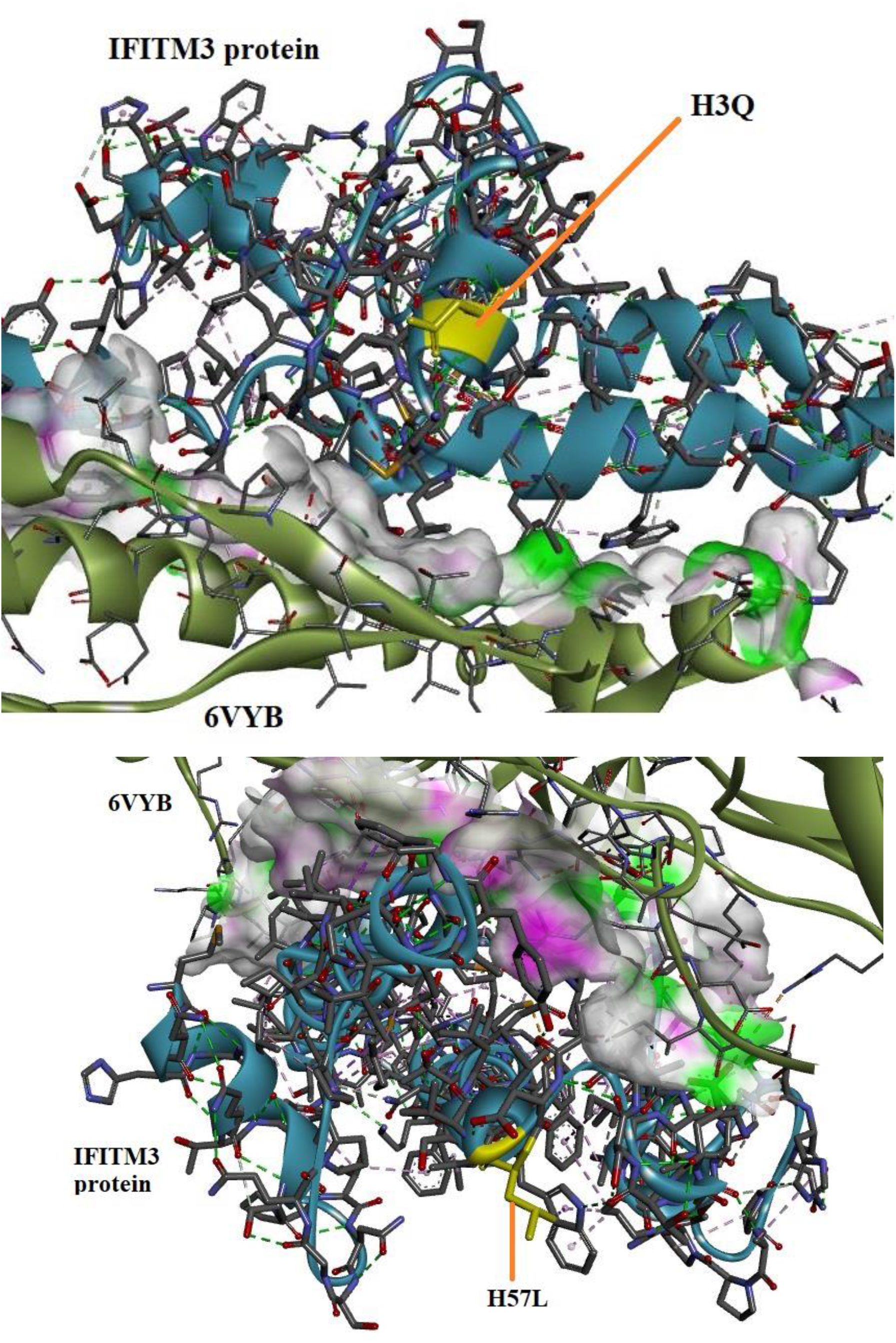
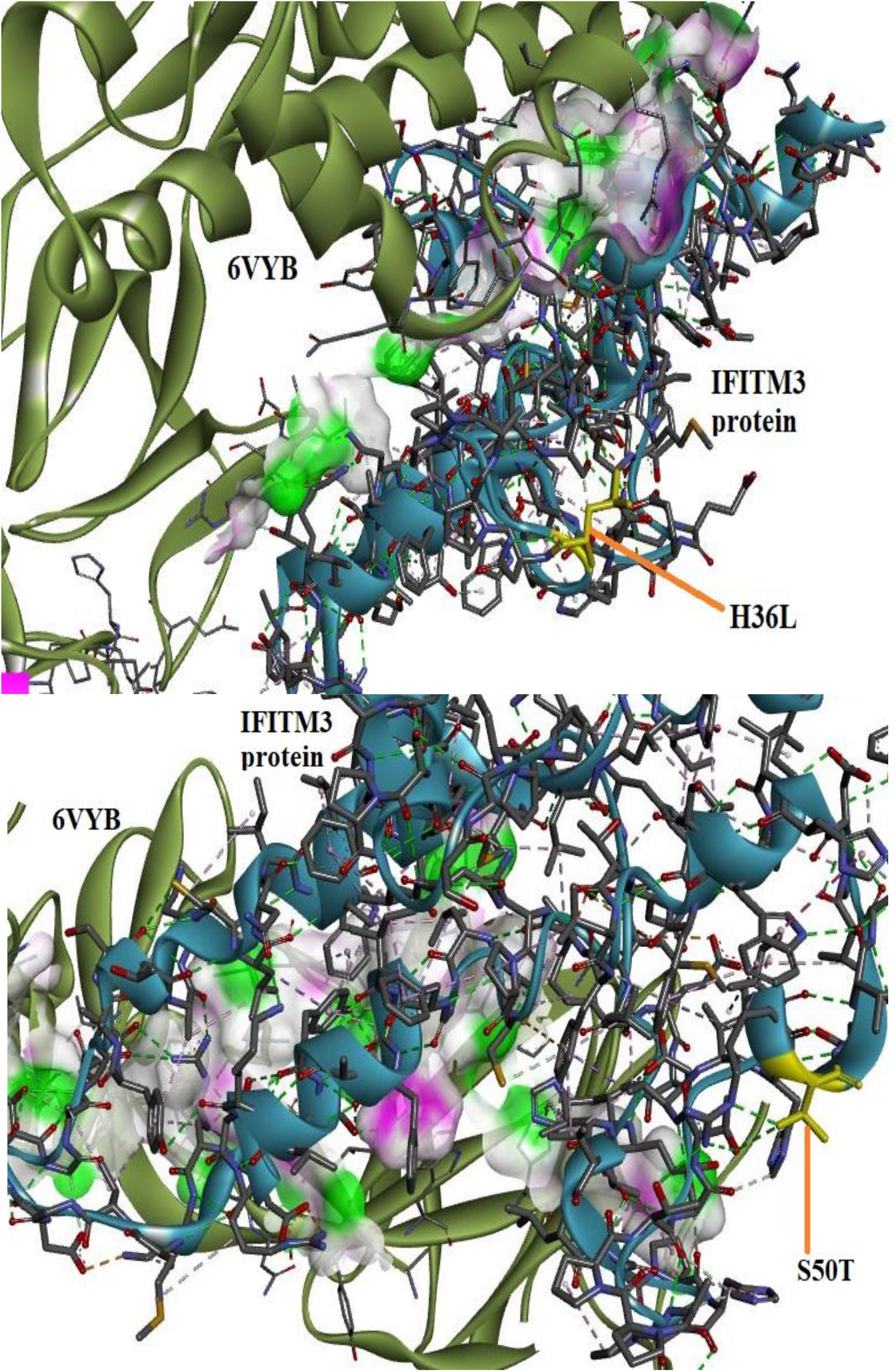
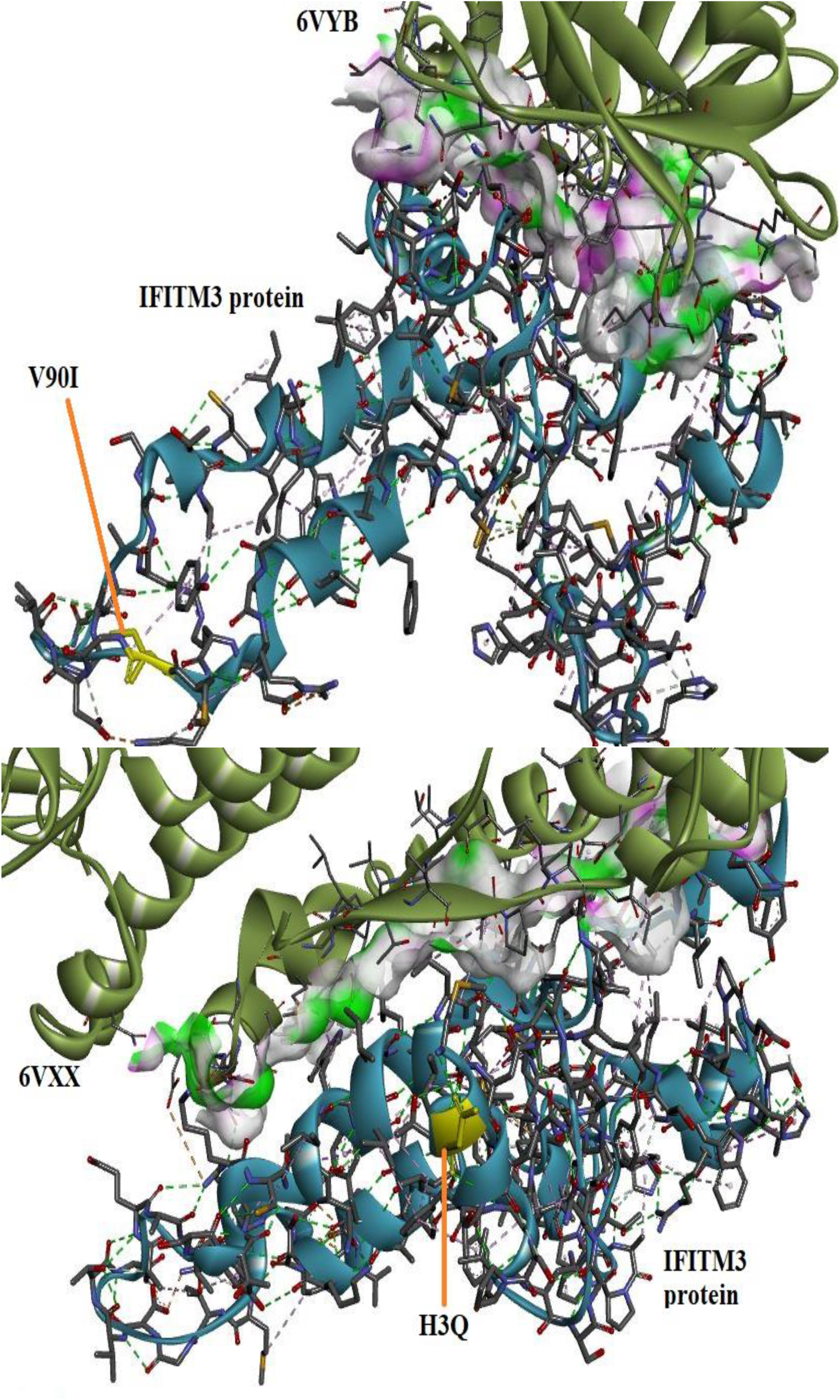
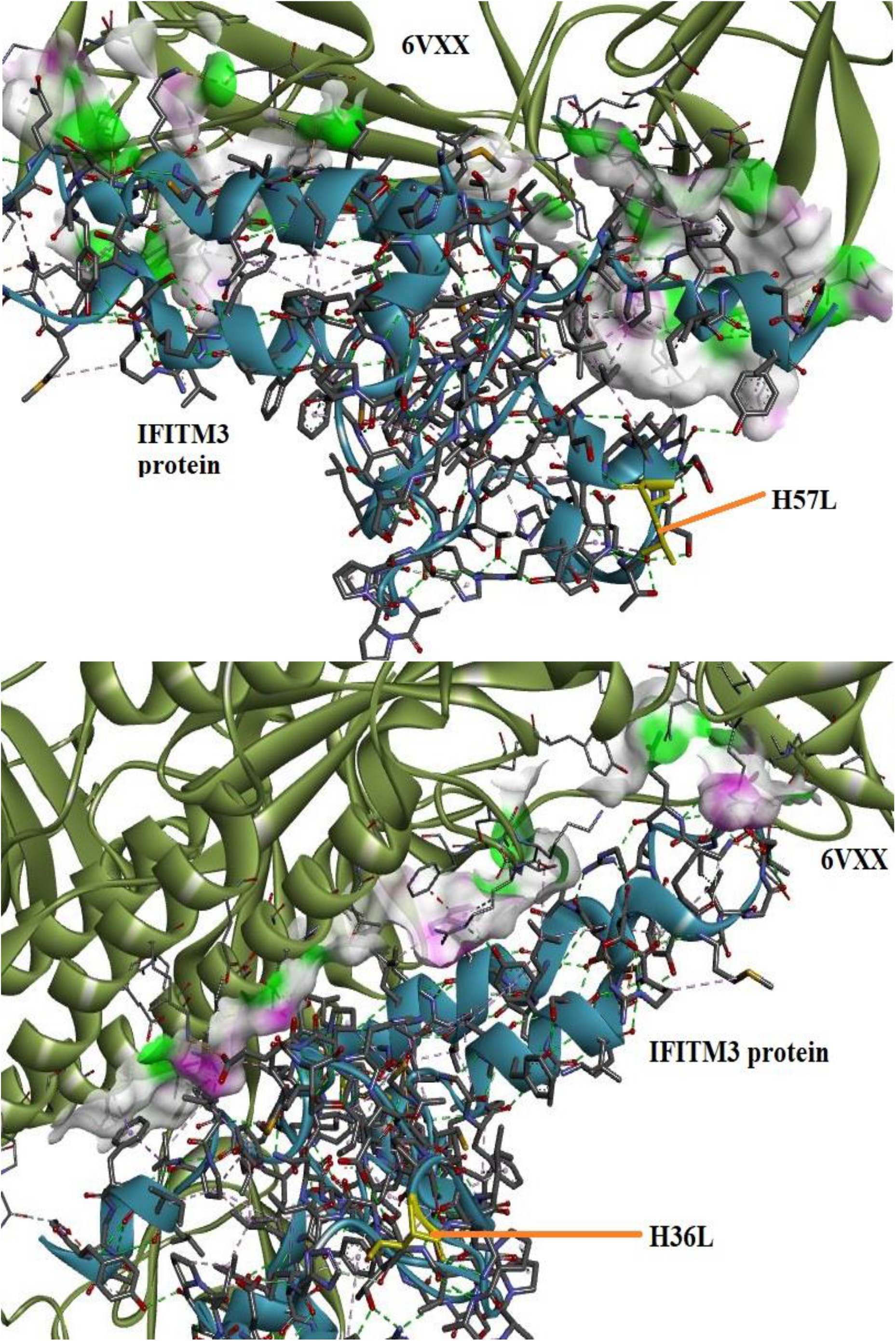
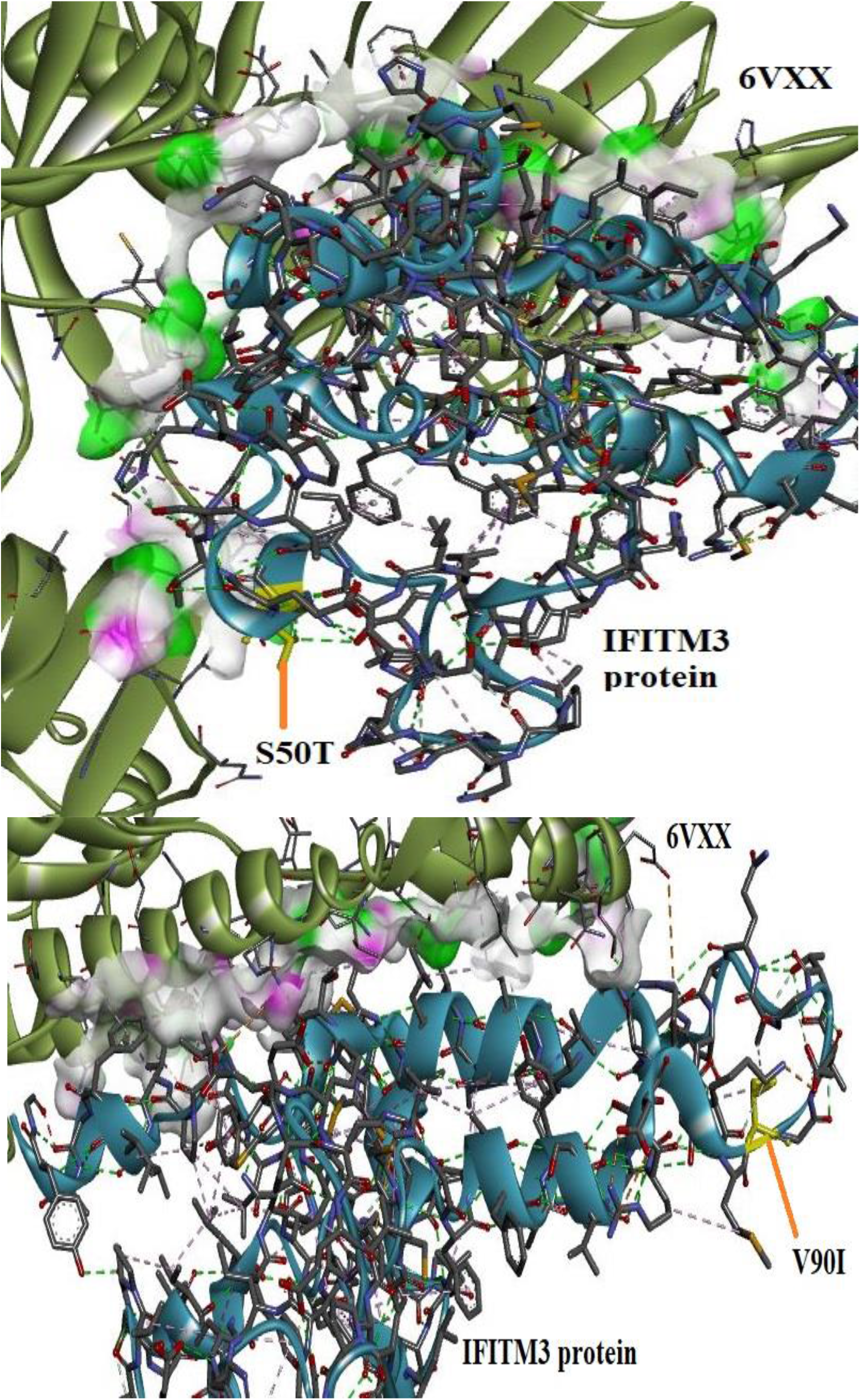
Visualization of the 6VYB/6VXX-IFITM1/2/3 (mutant) complexes via BIOVIA Discovery Studio.

## RESULTS AND DISCUSSION

### Data extraction and evaluation of the missense SNPs of IFITM1, IFITM2, and IFITM3

There were 1426 SNPs for *IFITM1*, 1834 SNPs for *IFITM2,* and 1584 SNPs for *IFITM3* in NCBI dbSNP on August 2021. Total missense SNP numbers for *IFITM1*, *IFITM2,* and *IFITM3* were 95, 139, and 130 respectively. rsIDs of the SNPs, allele changes, amino acid changes, pathogenicity/functional impact scores, predictions, functional impacts, and preservation times according to information from SIFT, PROVEAN, PolyPhen-2, SNAP2, Mutation Assessor, and PANTHER cSNP were shown in Table S1. 18 SNPs have harmful effects on the protein products of *IFITM1*, *IFITM2,* and *IFITM3* in at least four of the six tools. Among them, the most preserved are rs11552366 (*IFITM1*), rs200528039 (*IFITM1*), and rs371179996 (*IFITM2*) (456 million years). If the preservation is strong, the functional impact of the protein resulting from the variation will be high. Based on the information from whole tools, the most pathogenic missense SNP appears to be rs11552366 (*IFITM1*) (Table 1).

### DDG determination and structural analysis of the pathogenic forms of IFITM1, IFITM2, and IFITM3 proteins

According to the evaluation results of I-Mutant, MUpro, and SAAFEC-SEQ; amino acid changes in only two (H57L and H36L) of the 18 mutant proteins were predicted to increase the stability of the proteins (Table 2). Positive DDG values cause mutation to make the protein more stable, while negative values lead to more unstable proteins. The HOPE results showed that the mutant residue in seven (I87T, L41P, P34S, N63S, H3Q, H57L, and H36L) of the 18 mutant proteins had a smaller size compared to the wild-type residue; the mutant residues of the other 11 mutant proteins (Q120H, G74R, E30K, H6Y, V61M, V24M, C84Y, V53M, E20K, S50T, and V90I) were bigger than the wild-type residues. HOPE (Table 3) specified that changes in residue size and hydrophobicity due to mutant residue may affect contact with the lipid membrane. A mutant residue bigger than the wild-type residue may cause bumps. The results indicate that H6Y, N63S, H57L, H36L cause more hydrophobic protein structure. This is in agreement with the I-Mutant and MUpro results, as hydrophobicity can impart stability to mutant proteins. I87T, G74R, C84Y, and P34S lead to the formation of less hydrophobic protein structures, which can make the protein less stable. The reduced hydrophobicity of the protein means loss of H-bonds, resulting in loss of hydrophobic interactions, it also has a negative effect on protein folding. These results are also compatible with the protein structure analysis tools. Table x also shows that the charges of some mutant residues are different from the charges of wild-type residues. The difference in the charge of the mutant-type residue from that of the wild-type residue can cause repulsions within the protein itself or between the protein and the ligand.

### Homology modeling of wild and mutant types of IFITM1, IFITM2, and IFITM3 proteins

As a result of homology modeling evaluation for wild-type protein models provided by I-TASSER, the models with the highest C, ERRAT, and QMEAN scores were selected, therefore the homology models with the most quality are Model 1 for IFITM1, Model 1 for IFITM2, and Model 2 for IFITM3 (Table 4). The C and ERRAT scores appear to be within acceptable ranges, but the QMEAN scores are lower than expected in all models.

**Table 4.** Homology model validation of the IFITM1, IFITM2, and IFITM3 proteins

| **I-TASSER**<br>**Homology**<br>**models** | **IFITM1** |  |  | **IFITM2** |  |  | **IFITM3** |  |  |
| --- | --- | --- | --- | --- | --- | --- | --- | --- | --- |
|  | **C-score** | **QMEAN** | **ERRAT** | **C-score** | **QMEAN** | **ERRAT** | **C-score** | **QMEAN** | **ERRAT** |
| **Model 1** | -3.19 | -8.08 | 81.1966 | -4.14 | -14.08 | 81.4516 | -4.04 | -14.59 | 44.8 |
| **Model 2** | -3.60 | -9.43 | No result | -4.29 | -14.56 | 39.5161 | -4.42 | -11.39 | 77.6 |
| **Model 3** | -4.53 | -9.67 | 81.1966 | -4.78 | -11.79 | 58.871 | -4.24 | -14.63 | 61.6 |
| **Model 4** | -5 | -10.18 | 73.5043 | -5 | -14.48 | 58.871 | -4.35 | -13.78 | 47.2 |
| **Model 5** | -4.89 | -8.55 | 85.4701 | -4.46 | -14.18 | 44.3548 | -4.82 | -12.56 | 68 |

### Molecular docking of protein-protein complexes

As a result of the docking analysis performed with HawkDock (Table 5), the best binding among the open form of Spike-IFITM (wild) complexes belong to 6VYB-IFITM1 and the binding energy of the complex is -46.16 kcal/mol. This is followed by 6VYB-IFITM3 (-31.63 kcal/mol) and 6VYB-IFITM2 (-22.7 kcal/mol) complexes. The best binding among the close form of the spike (6VXX)-IFITM (wild) complexes belongs to 6VXX-IFITM1 (-52.42 kcal/mol). The binding energies of 6VXX-IFITM3 (wild) and 6VXX-IFITM2 (wild) complexes are -41.29 kcal/mol and -26.67 kcal/mol, respectively.

**Table 5.** Binding energies of the Spike-IFITM complexes

| COMPLEX |  |  |  | COMPLEX |  |  |  |
| --- | --- | --- | --- | --- | --- | --- | --- |
| PROTEIN | LIGAND | SCORE | BINDING FREE ENERGY (kcal/mol) | PROTEIN | LIGAND | SCORE | BINDING FREE ENERGY (kcal/mol) |
| 6VYB | IFITM1 (WILD) | -4947.78 | -46.16 | 6VXX | IFITM1 (P34S) | -5351.80 | -39.17 |
| 6VYB | IFITM1 (I87T) | -4193.10 | -29.14 | 6VYB | IFITM2 (WILD) | -5753.37 | -22.7 |
| 6VYB | IFITM1 (Q120H) | -4769.11 | -45.65 | 6VYB | IFITM2 (V53M) | -5899.53 | -22.43 |
| 6VYB | IFITM1 (G74R) | -4572.53 | -23.03 | 6VYB | IFITM2 (E20K) | -5729.59 | -22.35 |
| 6VYB | IFITM1 (E30K) | -4866.31 | -28.76 | 6VYB | IFITM2 (N63S) | -5567.96 | -25.37 |
| 6VYB | IFITM1 (H6Y) | -4868.30 | -46.28 | 6VXX | IFITM2 (WILD) | -4680.82 | -26.67 |
| 6VYB | IFITM1 (V61M) | -4371.18 | -33.43 | 6VXX | IFITM2 (V53M) | -5046.25 | -22.5 |
| 6VYB | IFITM1 (V24M) | -5154.42 | -38.57 | 6VXX | IFITM2 (E20K) | -5026.56 | -35.48 |
| 6VYB | IFITM1 (C84Y) | -4985.04 | -45.63 | 6VXX | IFITM2 (N63S) | -5169.55 | -50.77 |
| 6VYB | IFITM1 (L41P) | -4494.49 | -14.76 | 6VYB | IFITM3 (WILD) | -4507.06 | -31.63 |
| 6VYB | IFITM1 (P34S) | -4744.98 | -48.25 | 6VYB | IFITM3 (H3Q) | -4540.09 | -30.64 |
| 6VXX | IFITM1 (WILD) | -5477.52 | -52.42 | 6VYB | IFITM3 (H57L) | -4651.07 | -44.09 |
| 6VXX | IFITM1 (I87T) | -4961.61 | -31.89 | 6VYB | IFITM3 (H36L) | -4537.24 | -36.28 |
| 6VXX | IFITM1 (Q120H) | -4711.17 | -20.89 | 6VYB | IFITM3 (S50T) | -4808.95 | -22.31 |
| 6VXX | IFITM1 (G74R) | -4579.18 | -17.5 | 6VYB | IFITM3 (V90I) | -5012.83 | -42.91 |
| 6VXX | IFITM1 (E30K) | -4934.25 | -39.43 | 6VXX | IFITM3 (WILD) | -4757.20 | -41.29 |
| 6VXX | IFITM1 (H6Y) | -4948.96 | -36.16 | 6VXX | IFITM3 (H3Q) | -5048.03 | -33.15 |
| 6VXX | IFITM1 (V61M) | -5148.02 | -32.58 | 6VXX | IFITM3 (H57L) | -4766.14 | -31.07 |
| 6VXX | IFITM1 (V24M) | -5143.98 | -32.27 | 6VXX | IFITM3 (H36L) | -4563.16 | -14.48 |
| 6VXX | IFITM1 (C84Y) | -4993.99 | -31.37 | 6VXX | IFITM3 (S50T) | -4733.36 | 4.86 |
| 6VXX | IFITM1 (L41P) | -4526.06 | -47.08 | 6VXX | IFITM3 (V90I) | -5153.37 | -32.82 |

Bond analysis (H-bond, salt bridge, and non-contact bond) by PDBSum (Table 6 and Table 7) shows that the number of H-bonds between 6VYB and IFITM1/IFITM3/IFITM3 (wild) is equal (two H-bonds). Hydrogen bonds play an important role in determining the three-dimensional structures and the specificity of ligand binding [49, 50]. It is reported that H-bonds displace protein-bound water molecules and promote the ligand-binding affinity [51]. The 6VYB-IFITM3 (wild) complex differs from other 6VYB-IFITM (wild) complexes in that it has a salt bridge in between. In terms of non-bonded contact numbers, there are 158 bonds between 6VYB and IFITM2 (wild), while 150 bonds with IFITM3 (wild) and 131 bonds with IFITM3 (wild). Although the number of non-bonded contacts is less in the 6VYB-IFITM3 (wild) complex than other 6VYB-IFITM (wild) complexes, the extra salt bridge may explain the binding energy of the complex being better than the binding energy of the 6VYB-IFITM2 (wild) complex. The difference between the binding energies of the 6VXX-IFITM (wild) complexes emphasized the importance of a large number of non-bonded contacts. There are 197, 144 and 147 non-bonded contacts between protein-protein in the 6VXX-IFITM1 (wild), 6VXX-IFITM2 (wild) and 6VXX-IFITM3 (wild) complexes, respectively. Although the 6VXX-IFITM3 complex has six H-bonds and four salt bridges, the binding energy of the complex is less negative than the 6VXX-IFITM1 (wild) complex. Non-covalent interactions are relatively weak interactions, but their excess may provide a better interaction than H-bonds and salt bridges.

**Table 6.** Numbers of H-bonds, salt bridges and non-bonded contacts of the Spike-IFITM complexes

| COMPLEX |  |  |  |  | COMPLEX |  |  |  |  |
| --- | --- | --- | --- | --- | --- | --- | --- | --- | --- |
| PROTEIN | LIGAND | No of H-bonds | No of salt bridges | No of non-bonded contacts | PROTEIN | LIGAND | No of H-bonds | No of salt bridges | No of non-bonded contacts |
| 6VYB | IFITM1 (WILD) | 2 | - | 150 | 6VXX | IFITM1 (P34S) | 2 | 1 | 143 |
| 6VYB | IFITM1 (I87T) | 1 | 1 | 103 | 6VYB | IFITM2 (WILD) | 2 | - | 158 |
| 6VYB | IFITM1 (Q120H) | 2 | - | 152 | 6VYB | IFITM2 (V53M) | 2 | - | 160 |
| 6VYB | IFITM1 (G74R) | 3 | 1 | 154 | 6VYB | IFITM2 (E20K) | - | - | 220 |
| 6VYB | IFITM1 (E30K) | 5 | - | 147 | 6VYB | IFITM2 (N63S) | 2 | - | 159 |
| 6VYB | IFITM1 (H6Y) | 2 | - | 151 | 6VXX | IFITM2 (WILD) | - | 1 | 144 |
| 6VYB | IFITM1 (V61M) | 2 | 1 | 191 | 6VXX | IFITM2 (V53M) | 2 | - | 196 |
| 6VYB | IFITM1 (V24M) | 8 | 1 | 190 | 6VXX | IFITM2 (E20K) | 5 | - | 188 |
| 6VYB | IFITM1 (C84Y) | 2 | - | 152 | 6VXX | IFITM2 (N63S) | 4 | 1 | 148 |
| 6VYB | IFITM1 (L41P) | 7 | - | 141 | 6VYB | IFITM3 (WILD) | 2 | 1 | 131 |
| 6VYB | IFITM1 (P34S) | 2 | - | 148 | 6VYB | IFITM3 (H3Q) | 2 | 1 | 129 |
| 6VXX | IFITM1 (WILD) | 1 | - | 197 | 6VYB | IFITM3 (H57L) | 4 | 1 | 296 |
| 6VXX | IFITM1 (I87T) | 4 | - | 150 | 6VYB | IFITM3 (H36L) | 3 | 1 | 106 |
| 6VXX | IFITM1 (Q120H) | 4 | 1 | 142 | 6VYB | IFITM3 (S50T) | 2 | 1 | 223 |
| 6VXX | IFITM1 (G74R) | 4 | - | 150 | 6VYB | IFITM3 (V90I) | 4 | - | 155 |
| 6VXX | IFITM1 (E30K) | 3 | - | 223 | 6VXX | IFITM3 (WILD) | 6 | 4 | 147 |
| 6VXX | IFITM1 (H6Y) | 6 | 2 | 164 | 6VXX | IFITM3 (H3Q) | 3 | - | 149 |
| 6VXX | IFITM1 (V61M) | 3 | - | 174 | 6VXX | IFITM3 (H57L) | 1 | 1 | 210 |
| 6VXX | IFITM1 (V24M) | 4 | - | 150 | 6VXX | IFITM3 (H36L) | 2 | 3 | 148 |
| 6VXX | IFITM1 (C84Y) | - | - | 188 | 6VXX | IFITM3 (S50T) | 4 | 2 | 149 |
| 6VXX | IFITM1 (L41P) | 4 | 2 | 148 | 6VXX | IFITM3 (V90I) | 3 | 1 | 149 |

**Table 7.**
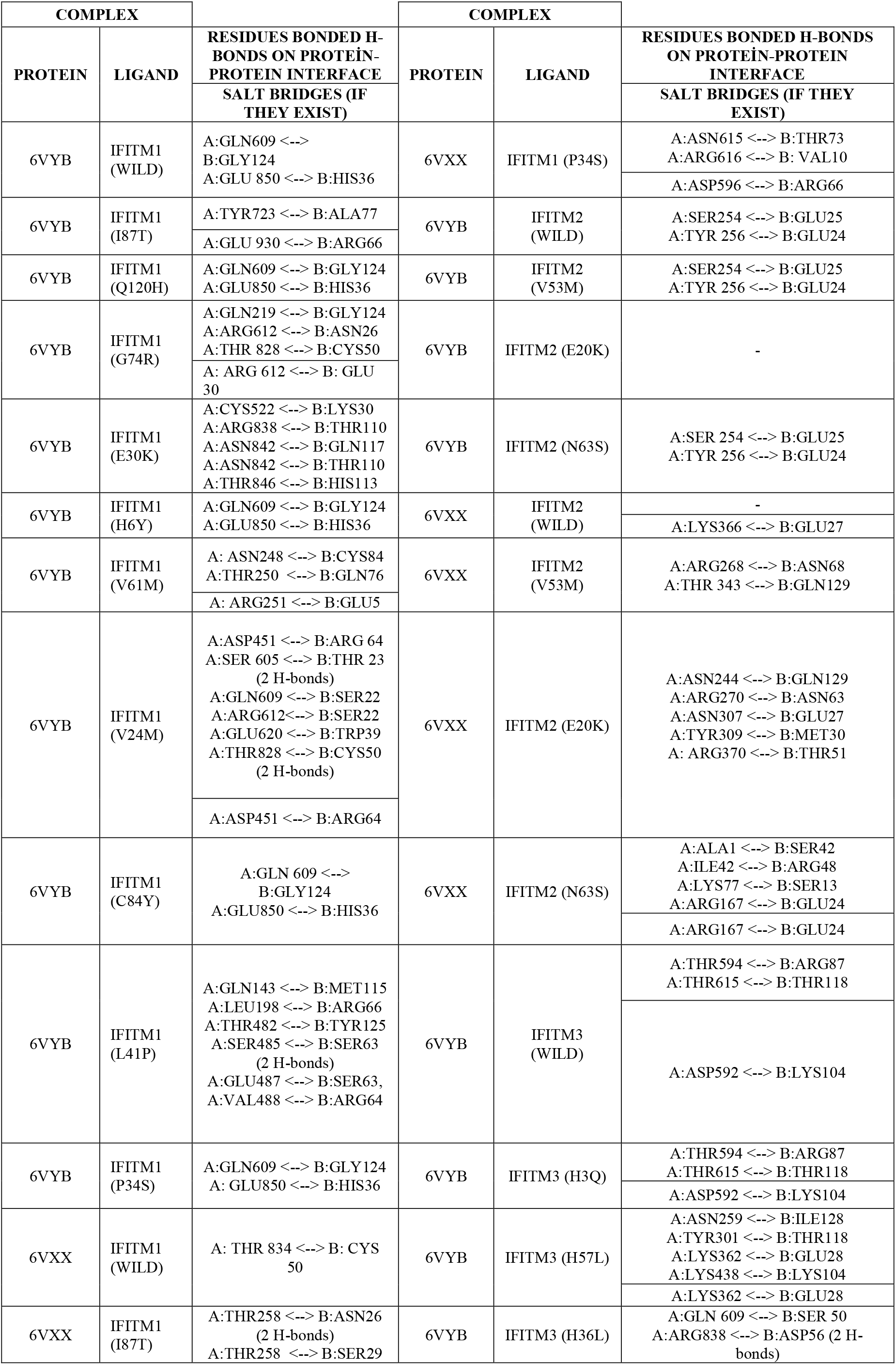
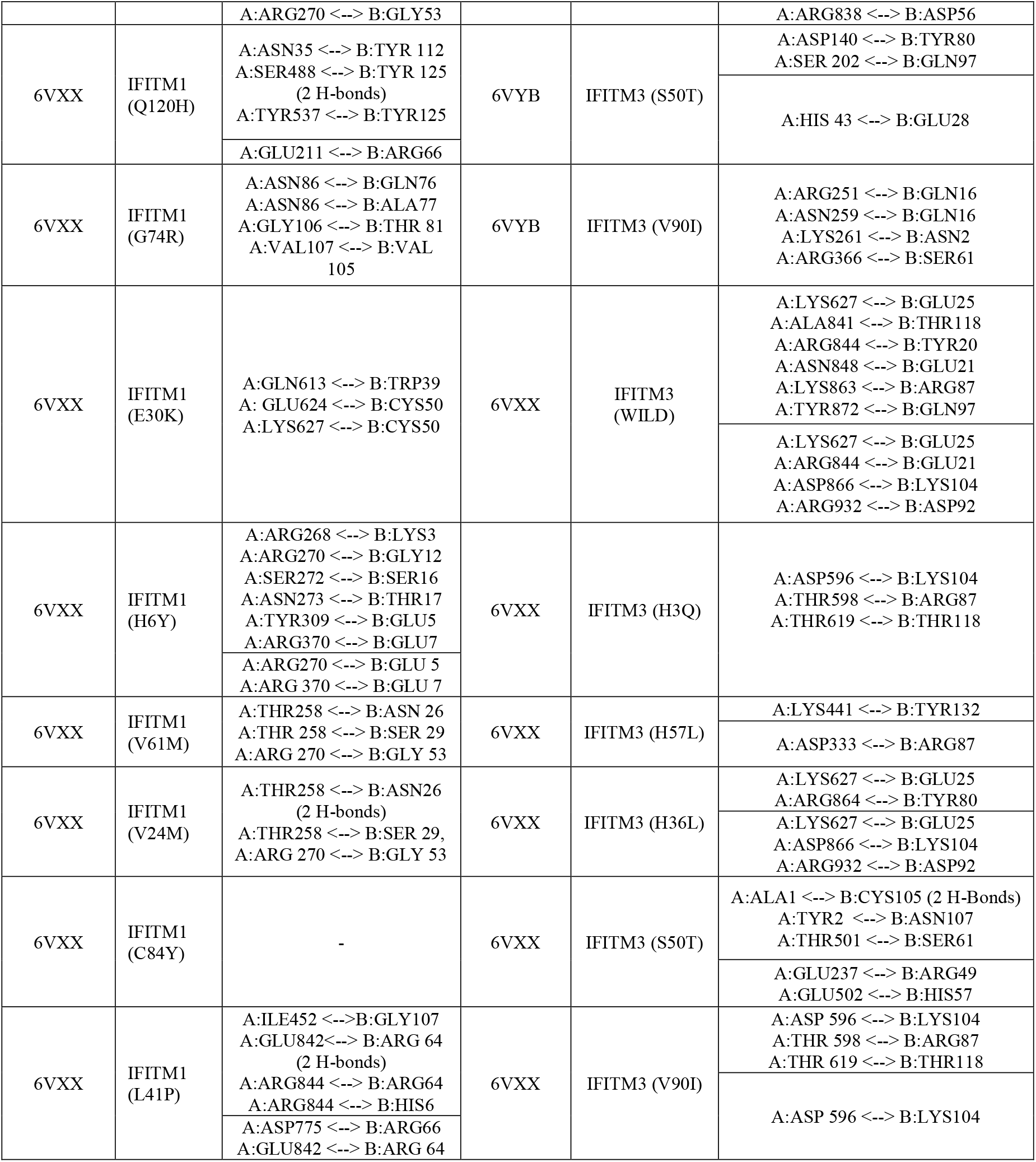
The positions of the H-bonds and the salt bridges of Spike-IFITM complexes

In Spike-IFITM (wild) complexes, the places where H-bonds are formed are different according to the closed and open forms of the Spike. H-bond formation with the S1 domain of the spike is seen in all 6VYB-IFITM (wild) complexes. Only 6VYB-IFITM1 (wild) complex also has H-bond formation of IFITM1 (wild) with S2. There is H-bond formation with both the S1 and S2 in the 6VXX-IFITM3 (wild) complex, while there is only H-bond formation with the S1 in the 6VXX-IFITM1 (wild) complex. There is no H-bond between 6VXX and IFITM2 (wild). Interesting results have emerged regarding the sites where H-bonds are formed, both with the 6VYB-IFITM (wild) and 6VXX-IFITM (wild) complexes. In the 6VYB-IFITM1 complex, binding occurred between the S1 and extracellular region of IFITM1 and between S2 and the cytoplasmic region of IFITM1. This may indicate that IFITM1 protein acts as a receptor for the Spike protein.

The most negative binding energy belonged to the 6VYB-IFITM1 P34S between the mutant type proteins of IFITM1 and the 6VYB complexes; and showed a little better binding than the wild type of IFITM1. This complex contains two H-bonds at the same positions as the 6VYB-IFITM1 (wild) complex and does not contain salt bridges. This complex contains H-bonds at the same positions as the 6VYB-IFITM1 (wild) complex and does not contain salt bridges. The binding between amino acid 850 (GLU) of S2 and amino acid 36 (HIS) of IFITM1 in one of the cytoplasmic domains in the region may be somewhat restricted by proline, amino acid 34 of wild-type IFITM. Because HOPE results indicated that the new residue (serine) was smaller at this location. In this case, it can be clarified why mutant-type binding is better. There is no 6VXX-IFITM1 (mutant) complex that has a better binding energy value than the 6VXX-IFITM1 (wild) complex.

The number of non-bonded contacts in all of the complexes formed by the mutant type protein of 6VYB and IFITM2 is higher than the non-bonded contacts in the 6VYB-IFITM2 (wild) complex, and therefore the binding energies of all complexes are slightly more negative than the binding energy of the 6VYB-IFITM2 (wild) complex. In the complexes formed by mutant-type proteins of IFITM2 with both 6VYB and 6VXX, the most negative binding energy was observed in the complexes with IFITM2 N63S. The binding energies of 6VYB-IFITM2 N63S and 6VXX-IFITM2 N63S complexes are -25.37 kcal/mol and -50 kcal/mol, respectively.

Among the complexes formed by 6VYB and IFITM3 mutants, the most negative binding energy belongs to the 6VYB-H57L complex (-44.09 kcal/mol). There may be two reasons for this. Compared to the 6VYB-IFITM3 (wild) complex, the number of H-bonds in the 6VYB-H57L complex is 2 times higher (4) and the number of non-bonded contacts is slightly more than double (296). Between the complexes formed by 6VXX and IFITM3 mutants, there is no complex with more negative binding energy than the 6VXX-IFITM3 (wild) complex. Between the complexes formed by 6VXX and IFITM3 mutants, there is no complex with more negative binding energy than the 6VXX-IFITM3 (wild) complex. The 6VXX-IFITM3 (wild) complex has more H-bonds and salt bridges than the 6VXX-IFITM3 mutant complex. An interesting result here is that the binding energy (4.86 kcal/mol) of the 6VXX-S50T complex is positive, making it the worst binding. Unlike other 6VXX-IFITM3 complexes, in the 6VXX-S50T complex, three H-bonds are formed with the first and second amino acids of the signal peptide located at the N-terminus of the Spike protein and the cytoplasmic domain of IFITM3 S50T. It is difficult to say whether this is related to any mechanistic event, since there may also be a binding affinity seen only *in silico*.

## CONCLUSION

Although effective vaccines have been developed for COVID-19, there are still points to be clarified about the mechanism of SARS-CoV-2. The mechanisms need to be well known in the development of new inhibitory drugs for the treatment of viral diseases. This study is the only study that offers a molecular docking approach to determine the interaction of both wild and mutant types of IFITM proteins with the Spike protein of SARS-CoV-2. With this study, we propose for the first time that there may be another receptor (IFITM1) for Spike other than ACE2, TMPRSS2 and FURIN. Our study is *in silico*-based and needs to be investigated in vitro as well.

